# Seeing the world in a grain of sand: Fine global and local representations in foveal and parafoveal vision

**DOI:** 10.64898/2026.06.16.732548

**Authors:** Jiapeng Yin, Zheyuan Chen, Yizhejun Li, Wenheng Xie, Weibin Song, Tianye Wang, Ye Liu, Yingfan Liu, Hetian Cao, Xiaotao Wang, Yongyu Wang, Lixuan Liu, Jinghao Ge, Xiaohong Li, Lothar Spillmann, Stewart Shipp, Niall McLoughlin, Ian Max Andolina, Yiliang Lu, Shiming Tang, Wei Wang

## Abstract

Foveal vision enables primates to perceive small objects with remarkable precision by resolving both fine-global form and fine-local detail. Yet current coarse-to-fine and local-to-global frameworks, derived largely from parafoveal recordings and large-object representations, posit that V1 encodes local detail whereas IT represents global form, leaving the neuronal basis of foveal object representation unresolved. Here we show that fine-global form and fine-local detail are encoded parallelly as early as V1 and preserved across V2, V4 and IT at both foveal (1°–2°) and parafoveal (2°–6°) retinal eccentricities. Coarse-global features dominate parafoveal vision and higher cortical areas. Response latencies revealed a rapid processing sequence, with 2–6 ms separating coarse-global, fine-local, and fine-global processing within parafoveal V1, V2 and V4. These findings establish a cortical neuronal framework to resolve fine-global form of small objects in foveal vision along primate ventral visual stream, supporting fine-scale perception during reading and fine object recognition.

Highlights

1. Foveal two-photon imaging across V1, V2, V4, and IT
2. Global and local processing examined across eccentricity
3. Fine and coarse global representations emerge as early as V1
4. Fine global and local representations are preserved from V1 up to IT
5. Tightly timed coarse-to-fine-to-fine processing within each cortex
6. Fast coarse-global dominates parafoveal and higher cortical areas

## Introduction

Visual perception has fascinated scientists and philosophers for centuries. What appears effortless to us is the perception of objects in terms of both their global forms and local parts, with remarkable precision in the fovea. The fovea is a small, capillary-free depression at the center of the macula, approximately 0.4–0.5 mm in diameter corresponding approximately to the central 2° of retinal eccentricity. It contains the highest concentration of cone photoreceptors, which exhibit relatively slow temporal responses and high-fidelity signaling specialized for high-acuity form vision^1–3^ (**Figure 1A**). This is reflected in visual acuity testing, in which both small and large optotypes, such as the letter “E,” can be seen clearly when presented in the fovea. However, the cortical mechanisms underlying the joint representation of the global form and local parts of a coherent object—for example, the letter “E” and its constituent strokes—remain unclear, especially in foveal vision, where global form and local detail are perceived with comparable sharpness. This question, foreshadowed by William Blake’s poetic musings, gained scientific substance with the work of the Gestalt psychologists. They studied the psychophysical laws of visual perception and proposed that the global percept of the whole is processed prior to the local parts of a stimulus^4,5^, emphasizing that the whole has emergent properties that are not the simple sum of its parts. This perspective laid the conceptual foundation for subsequent theoretical and physiological work on local–global representations in the visual brain.

**Figure 1.**
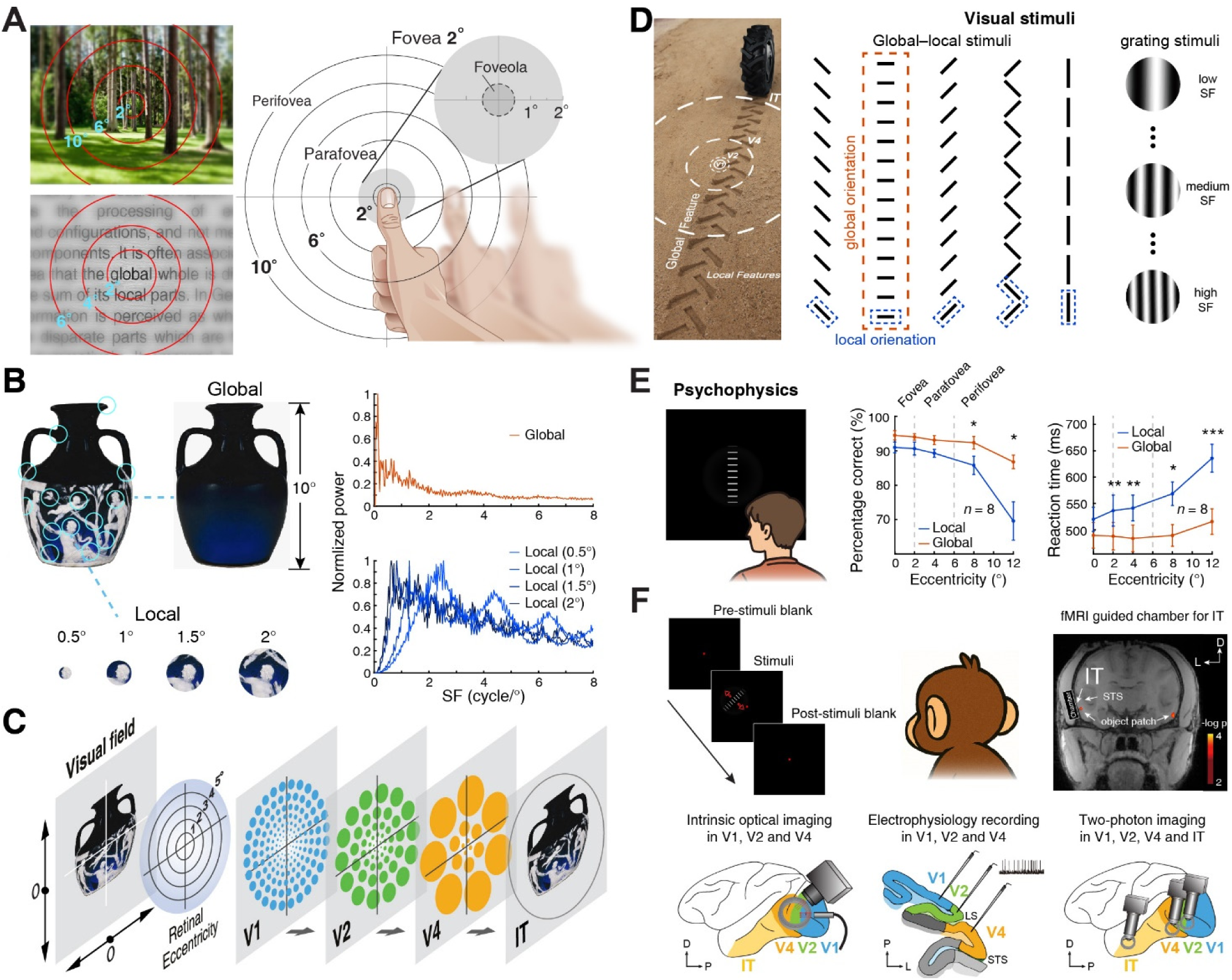
Distinct perception of global and local features in the fovea and beyond. **(A)** Comparisons of foveal and extrafoveal vision. In foveal vision, both local and global features in fine details are seen clearly, whereas in parafovea and beyond only the gist of the coarse global is perceived. The thumb illustrates distinct visual perception across retinal eccentricity at an arm distance. **(B)** SF power spectra of global and local features in the Portland Vase. When viewed from ∼1.5 m, the global feature subtends approximately 10°, and local image regions were sampled with apertures ranging from 0.5° to 2°. As the aperture increased, the SF power spectra of the image shifted systematically toward lower SFs. **(C)** The central question of this study: cortical cellular mechanisms underlying the foveal representations of fine global and local features across the primate ventral visual pathway. **(D)** Experimental stimuli design. Left panel, a photo of tire tracts in which global and local features differ drastically in spatial scales and orientations. Circles of different sizes superimposed on the photo indicate the approximate RF sizes of V1, V2, V4 and IT neurons. Right panel, the visual stimuli in this study (see Methods). **(E)** Psychophysical validation experiments with global–local stimuli (see Methods). *n* indicates the number of participants. \**p* < 0.05; \*\**p* < 0.01; \*\*\**p* < 0.001 with paired t-test. Dots and error bars represent the mean and standard error of the mean (SEM). **(F)** Neural recordings. fMRI-guided neuronal and population recordings across retinal eccentricity at foveal and parafoveal regions of each cortex from V1 to IT of awake macaques.

Although global and local features have distinct visual characteristics, both can be characterized by distance-dependent spatial-frequency (SF) composition, which can be quantified using classical Fourier analysis of their spatial power spectra^6^. For example, Fourier analysis of the Portland Vase image revealed that local features are biased toward higher-SF components, whereas global structure is dominated by lower SFs (**Figure 1B**). Notably, the power spectrum shifts toward lower SFs as the aperture size increases. Thus, low-SF visual information is classically assumed to convey the “gist” global structure^7–9^, and to precede high-SF local information in the primate ventral visual pathway^9–13^. This is referred to as coarse-to-fine analysis based on SF information analysis^9–11,13,14^, and is supported by neural recordings of parafoveal regions in V1, V2, and V4 toward object-selective regions in inferotemporal (IT) cortex^6,15–19^, where preferred SFs generally decrease along the ventral visual hierarchy when tested with sinusoidal gratings.

However, global shape is not necessarily defined only by low-SF information. For both small and large objects, their global shapes are defined mostly by sharp edges and lines, and natural objects and scenes often contain a broad range of low-and high-SF components^20,21^. Furthermore, neurons representing the foveal region have very small receptive fields (RFs) and support exceptionally high visual acuity^15–17^. This high acuity allows foveal vision to resolve not only local details but also emergent global forms defined by high-SF components (**Figure 1AB**). Accordingly, as visual acuity declines exponentially with increasing eccentricity, global form remains readily perceived but becomes blurred in the parafovea and peripheral vision (2–6° and beyond)^22–24^. Furthermore, cortical neuronal recordings from parafoveal regions have revealed that the sizes of neuronal RFs increase markedly along the primate ventral visual pathway^25–28^, allowing cortical neurons to integrate information over progressively larger regions of visual space and to represent large objects and scenes of increasing complexity^26–30^. These findings form the basis of a dominant hierarchical feature-integration framework, which posits that V1 primarily encodes local details whereas IT represents objects and scenes through progressive pooling and integration of lower-level features. However, this framework, which emphasizes a transformation from local to global representations, does not specify how fine global form of small objects is jointly represented with fine local information within foveal representations.

As neuronal RF size increases with retinal eccentricity and along the visual hierarchy^15–17,27,31^, while visual acuity decreases radically, the spatial contents available for cortical processing change substantially across both visual space and cortical stages. Thus, it remains unclear how global forms spanning a wide range of SFs, together with fine local features, are neuronally represented across foveal and parafoveal regions within each cortical area from V1 to IT (**Figure 1C**). This remains a central question for understanding cortical processing underlying fine-scale visual behaviors, including reading and foveal object recognition (**Figure 1A**).

## Results

The complexity of natural images, which contain multiple orientations and other visual attributes, limits their utility for isolating the neural mechanisms of global and local processing without applying oriented spatial filters^10,13,14,32^. To overcome these limitations, we constructed two-dimensional tire-track-like global–local stimuli composed of aligned, equally spaced local line elements (**Figure 1D**), which can be reliably identified in neuronal responses at both single-cell and population levels. The SFs of these global–local stimuli were set at 0.5 cycles/° and 3 cycles/° representing coarse global and fine local information, respectively. Conceptually similar to the hierarchical stimuli introduced by Navon^7^, these stimuli enabled psychophysical validation showing that global form and local features with orthogonal orientations are perceived with comparable precision within the central 2° of the fovea (**Figure 1E**). In contrast, global form becomes increasingly dominant in the parafoveal region and beyond. This eccentricity-dependent decline in acuity affects local feature perception more strongly than global structure, indicating that global and local processing vary across retinal eccentricity along cortical hierarchy.

To address cortical neuronal mechanisms underlying fine global and local representations in foveal and parafoveal vision, we performed two-photon imaging with single-cell resolution in awake macaques^33,34^, across four successive cortical areas along the ventral stream—V1, V2, V4, and IT—at eccentricities of 1–2° within the foveal region (**Figure 1F**). In the parafoveal regions of these cortices (2–6° eccentricity), we combined large-window intrinsic-signal optical imaging of population responses with extracellular electrophysiological recordings^6,34,35^.

### Distinct spatial components of global form and local features

In contrast to control stimuli of oriented sinusoidal gratings, whose spatial power is concentrated at a single SF, global–local stimuli contain multiple SF components (**Figure S1A**), consistent with the Fourier analysis results obtained from the image of the Portland Vase. Notably, in addition to spectral power being concentrated at the low SF that matches the designed global SF of 0.5 cycles/°, global forms also contain high-SF components, including those derived from the oriented line endpoints. Accordingly, the power spectrum of global form information spanned from coarse to fine (0.5 to 8 cycles/°). In contrast, fine local information remains sharp, as it is dominated by spectral power of high SFs, at the preset 3 cycles/° and above.

Importantly, as stimulus size increases progressively to match neuronal RFs across retinal eccentricity and individual cortical area along the visual hierarchy^27,28^, the proportion of global SF power rises rapidly relative to local SF power (**Figure S1BC**), suggesting that neurons with larger RFs are exposed to proportionally much more global information containing both low-and high-SF components. These results indicate that global form and local parts are represented in parallel but not in fixed proportion within inherently overlapping spatial-frequency domains. Note that stimulus scale and RF size increase across retinal eccentricity and the visual hierarchy, low-SF components increase markedly and dominate global representations. Thus, global and local representations arise from scale-dependent mixtures of low-and high-SF components rather than from strictly separable spatial-frequency bands. This spatial analysis challenges the common practice of using low-SF filtering as a proxy for global information^9,32^.

### Hierarchical population representations of global and local in the parafovea

Based on the spatial analysis of global form and local features (**Figure S1**), we first investigated how global and local information are represented at the population level in the parafoveal regions across the visual hierarchy. Cortical population responses in V1, V2, and V4 at eccentricities spanning from 1° to 6° were simultaneously recorded with intrinsic-signal optical imaging in eight hemispheres from five macaques (**Figure 1F**). Differential orientation maps (dark–bright domains) in response to drifting global–local stimuli, with angular differences between global and local orientations of 90°, 45°, and 0°, were obtained (**Figure 2A**). Oriented drifting sinusoidal gratings served as control stimuli. By analyzing orientation preference maps (colored contours) overlaid on the differential orientation maps for each area^6,35^ (**Figure S2**), we found that the population response profiles to grating stimuli in all visual areas exhibited peaks and troughs at the expected orientations (0°, 90°, 45° and 135°). For global–local stimuli, population responses in V1 and V2 aligned consistently with the local orientations. In contrast, those in V4 tracked only the global orientation and were unaffected by the orientations of local elements. To further illustrate global and local representations across retinal eccentricity in V1, V2, and V4, we constructed global–local preference maps by extracting the peak orientation of the response profile at each pixel in response to the above global–local stimuli (**Figure 2B**). In V1 and V2, peak orientations were consistently centered at around 90° or 45°, corresponding to the local orientations of the stimuli, whereas in V4 peak orientations only represented the global orientations. These hierarchical population responses—global-dominant in V4 and local-dominant in V1 and V2—were consistently observed across all hemispheres and animals (**Figures S3 and S4**).

**Figure 2.**
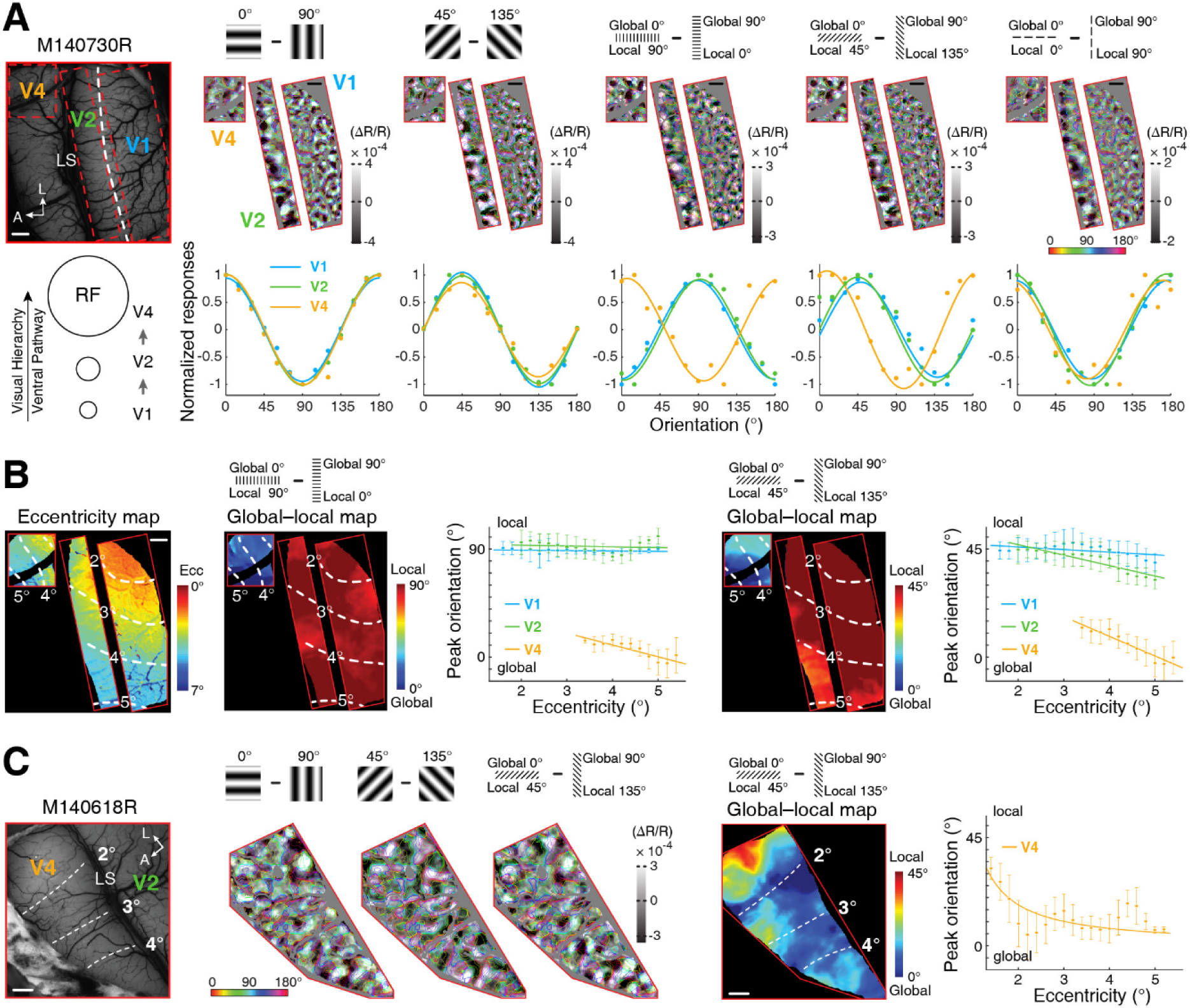
Population representations of global and local in the parafovea. **(A)** Hierarchical representations of global and local orientations. The leftmost column depicts the vasculature surface of areas V1, V2, and V4 in the right hemisphere of a macaque (M140730). Regions of interest (ROIs) outlined in red. The dashed line indicates the V1–V2 border defined by ocular dominance responses. Note that the RF size in V4 is at least four times larger than in V1^17,28^. A, anterior; L, lateral; LS, lunate sulcus. Scale bar, 1 mm. Differential orientation maps, generated by paired stimuli as shown in the stimulus icons, overlaid with colored iso-orientation contours derived from orientation preference maps. Orientation response profiles for V1, V2, and V4 are shown underneath their corresponding orientation maps. **(B)** Global–local preference maps across retinal eccentricity. The leftmost panel shows the eccentricity maps of V1, V2, and V4 from the same ROIs as in (A). The global and local preference maps for two paired global–local stimuli are shown, respectively, with eccentricity lines overlaid. The orientation–eccentricity relationship, with peak orientation of the above response profile at each pixel was plotted against retinal eccentricity, is shown in the right. Dots and error bars represent the mean and standard deviation. **(C)** Global and local orientation representation in the foveal and parafoveal regions of V4. The leftmost panel shows the cortical surface of V4 across eccentricities from 1° to 6° in macaque (M140618). Differential orientation and the global–local preference maps are both presented. The orientation–eccentricity relationship is also plotted.

Thus, at the population level in parafoveal regions, V1 and V2 predominantly encode local orientations, whereas V4 encodes global orientations in an invariant manner, consistent with previous studies using abutting-line illusory contours^35,36^.

However, we previously demonstrated that there exist high-SF functional domains in parafoveal V4 encoding local orientations of the global–local stimuli^6,37^, suggesting that both local and global features are represented in parallel in V4. To further investigate how global and local representations vary with retinal eccentricity, we focused on V4, a key intermediate stage linking early visual cortex with higher cortical areas. Because V4 has larger receptive fields and a lower cortical magnification factor than V1 and V2, a smaller cortical surface area in V4 can represent a relatively larger region of visual space. Within the global–local preference maps of peak responses (global: 0°, local: 45°), we observed red-hot and dark-blue regions encoding local and global orientations, respectively (**Figure 2C**). Importantly, local orientations dominated V4 in its foveal regions within 2° eccentricity, whereas at more peripheral regions peak orientations shifted toward global encoding. These results indicate that V4 retains fine local information from foveal V1 and V2 inputs, while increasingly representing global information across retinal eccentricity within its larger RFs. Due to the limited spatial resolution of intrinsic-signal optical imaging, however, it remains unclear how global form and local information are encoded neuronally at each cortex of the visual hierarchy.

### Neuronal encoding of global and local information within the parafovea

To address the above question at the neuronal level, we extracellularly recorded 155, 90, and 189 orientation-selective neurons in V1, V2, and V4, respectively, at eccentricities between 2° and 6° in three awake macaques (**Figure 1F**). We characterized the response tuning of individual neurons to global–local stimuli, in which global and local orientations differed by 90°. Oriented sinusoidal gratings were used as control stimuli to identify each neuron’s preferred orientation. Based on feature selectivity quantified by a global–local index (GLI)^6,35^, neurons were then classified as local-orientation-selective, dual-orientation-selective or global-orientation-selective (**Figure 3A and Figure S5A**). Note that a subset of neurons displayed dual tuning, with two response peaks corresponding to local and global orientations of global–local stimuli, respectively. These three types of neurons were found across all cortical areas, indicating that both global and local information from the same stimuli emerge at the neuronal level as early as V1 and propagate in parallel along the ventral visual hierarchy.

**Figure 3.**
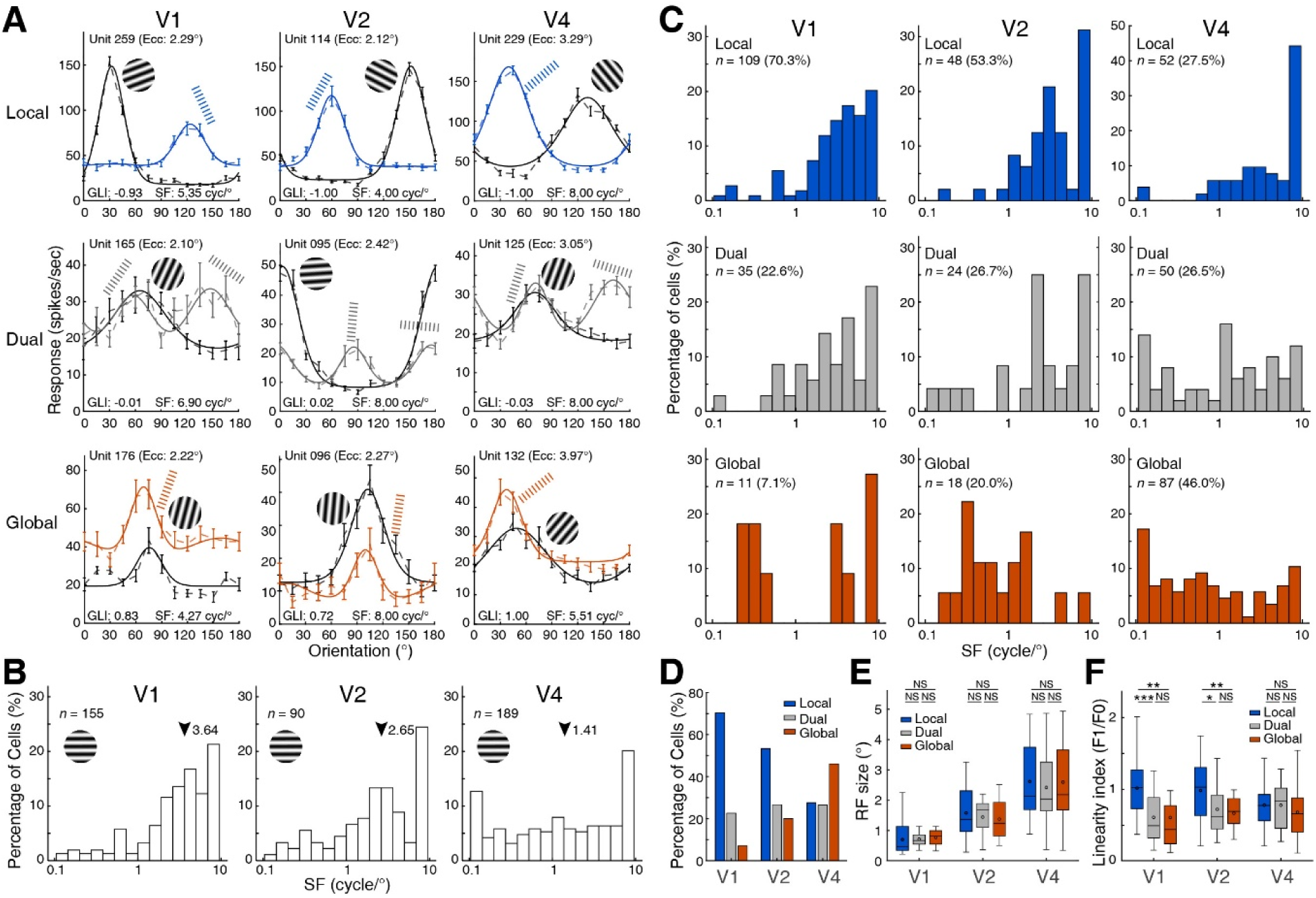
Neuronal encoding of global and local in the parafovea. **(A)** Examples of local-, dual-, and global-selective neurons in response to global–local stimuli in V1, V2, and V4. Stimulus icons indicate the preferred stimulus for each neuron. The x-axis labels denote grating orientations and global orientations in the global–local stimuli. GLI, global–local index; SF, neuron preferred SF measured with sinusoidal grating stimuli. Further global-selective neurons with low-SF preferences are presented in Figure S5A. **(B)** SF distributions of neurons sampled at each cortex, measured with sinusoidal gratings. Arrow indicates median SFs for each cortex. *n* indicates the number of neurons. **(C)** Distributions of the three neuron types, plotted against individual neuron’s preferred SF measured with sinusoidal grating stimuli. *n* indicates the number of neurons. **(D)** Proportions of local-, dual-, and global-selective neurons in each cortical area. **(E)** Comparison of RF sizes among local-, dual-, and global-selective neurons. Data are reported as mean ± SEM. For local-, dual-and global-selective neurons were 0.70 ± 0.06°, 0.72 ± 0.06° and 0.77 ± 0.17° in V1, 1.57 ± 0.16°, 1.44 ± 0.17° and 1.37 ± 0.42° in V2, and 2.62 ± 0.19°, 2.42 ± 0.18° and 2.59 ± 0.15° in V4, respectively. NS, not significant. **(F)** Comparison of the linearity indices among local-, dual-, and global-selective neurons. Post hoc Tukey–Kramer correction: \**p* < 0.05; \*\**p* < 0.01; \*\*\**p* < 0.001; NS, not significant.

In order to investigate underlying mechanisms responsible for these different types of local and global processing, we further examined their SF tunings using oriented sinusoidal grating stimuli with SFs ranging from 0.125 to 8 cycles/°. Consistent with previous findings^6,15–18^, cortical preferred SFs decreased along the visual hierarchy (median SFs: 3.64 cycles/°, 2.65 cycles/°, 1.41 cycles/° across V1, V2 and V4; one-way ANOVA, F = 7.00, *p =* 1.02 × 10⁻^3^) (**Figure 3B**). Note that the median preferred SF in V4 was higher than the ∼1 cycle/° reported in a previous study^17^. This is likely because many of our V4 recordings were obtained from high-SF domains as previously identified^6^.

Importantly, when plotted against their preferred SFs measured with sinusoidal gratings (**Figure 3C**), local-selective neurons across all areas were biased toward high SFs. By contrast, global-selective neurons spanned a broad range of preferred SFs, from 0.125 to 8 cycles/° (**Figure 3ABC and Figure S5A**), consistent with the spatial power analysis of the global–local stimuli (**Figure S1AB**). These results indicate that both coarse and fine global information can be successfully captured by neurons preferring low and high SFs, respectively, at each cortical area. Notably, a subset of neurons with low-SF preferences defined by sinusoidal grating stimuli (< 1 cycle/°; e.g., 12/109 in V1), responded well to the local orientation with its SF preset at 3 cycles/° (**Figure 3C and Figure S5**). This discrepancy is likely due to the fact that these low-SF preference neurons are broadband tuned (**Figure S5B**), although they showed significantly stronger responses at 0.5 cycles/° than to 3 cycles/° to sinusoidal grating stimuli. They were actually activated by the high-SF local information at 3 cycles/° which dominate the Fourier power spectrum of the global–local stimuli (**Figures S5C**).

Overall, at the neuronal level, global form emerges alongside local features as early as V1, but global dominance increases progressively from V1 to V4 (**Figure 3D**), consistent with population results obtained using intrinsic-signal optical imaging (**Figure 2 and Figures S2 and S3**). This gradual shift in the relative dominance of global and local encoding may be explained by increasing RF sizes across cortical areas (two-way ANOVA, F = 73.81, *p <* 0.001) (**Figure 3E**). By contrast, there was no significant difference in the RF sizes among the three cell types within each cortical area (two-way ANOVA, F = 0.40, *p =* 0.67), suggesting that RF size alone does not distinguish global from local representations within a given cortical area. To further distinguish global-and local-selective neurons, we compared the linearity of their responses to sinusoidal grating stimuli. In contrast to the linearity of local-selective neurons, both dual-selective and global-selective neurons exhibited greater nonlinearity in V1 and V2 (**Figure 3F**; one-way ANOVA, F = 19.31, *p <* 0.001 in V1, F = 7.34, *p =* 1.14 × 10⁻^3^ in V2, and F = 2.20, *p =* 0.12 in V4). Note that in V4 all three types including local-selective neurons exhibited nonlinearity. This nonlinearity suggests that the responses of dual-and global-selective neurons cannot be explained solely by the Fourier power spectrum of global–local stimuli, and may involve nonlinear spatial integration mechanisms, possibly including end-stopping as early as V1^25,36^. Thus, this stronger nonlinearity associated with global processing in V1 and V2 may reflect the emergent properties of global forms, consistent with the Gestalt view that the whole is not simply the sum of its parts.

Together, these parafoveal electrophysiological results reveal that both fine and coarse global information together with fine local information are encoded in parallel as early as V1, and across the cortices of the ventral visual hierarchy.

### Temporal processing of global and local information in the parafovea

The perceptual coarse (global) to fine (local) framework, in which global information is proposed to be processed before local detail, is supported by observations that neurons preferring low SFs exhibit shorter response latencies than those preferring higher SFs when tested with sinusoidal gratings^9,38^. These slower high-SF responses likely originate in the primate fovea, which is characterized by a high density of cone photoreceptors and midget ganglion cells with relatively slower temporal responses than those in the perifovea^2,39^. We confirmed the latency differences between low and high SF by using sinusoidal gratings in V1, V2 and V4 (**Figure 4AB**) and V4 in previous study^6^. In addition, these response latencies of either onsets or peaks exhibited monotonical increases for all SFs across the visual hierarchy. However, whether neuronal responses to coarse and fine global forms defined by low and high SFs, respectively, together with fine local detail within global–local stimuli are themselves temporally distinguishable has not been examined directly. Given the band-pass nature of SF tunings of neuronal responses to sinusoidal gratings^15–17^ and the broadband SF power spectrum of global information in the global–local stimuli (**Figure S1**), neurons preferring intermediate SFs (1–3 cycles/°) could not be assigned to either low-or high-SF global groups. Thus, we only compared response latencies of global neurons in response to global–local stimuli, but based on their preferred SFs were either below 1 cycle/° (low-SF global) or above 3 cycles/° (high-SF global) measured with sinusoidal gratings. As the SF components of local features in the global–local stimuli were 3 cycles/° and above (**Figure S1**), neurons in responses to local orientation were classified as high-SF local orientation selective. Therefore, responses latencies were compared only among low-SF global, high-SF global and high-SF local groups at each cortical area along the visual hierarchy (**Figure 4CD**).

**Figure 4.**
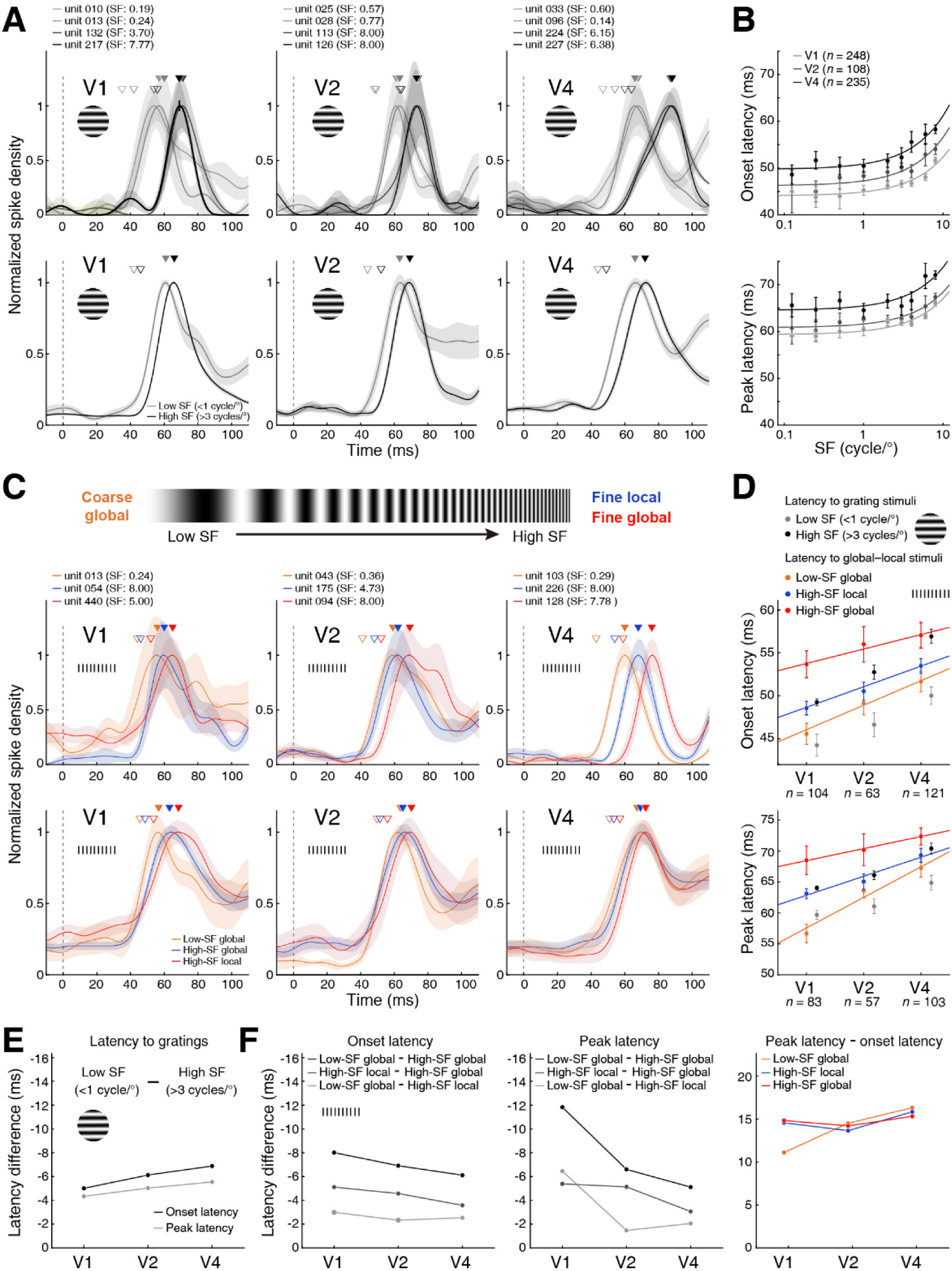
Response latencies to sinusoidal gratings and global–local stimuli from V1 to V4. **(A)** Temporal profiles of neuronal responses to gratings at each neuron’s preferred orientation and preferred SF. The top and bottom rows show responses from example neurons and population averages, respectively. Hollow and solid triangles indicate onset and peak latencies. **(B)** Distribution of onset and peak response latencies for sinusoidal grating stimuli. Latencies were measured from neuronal responses to the preferred orientation and preferred SF of each neuron. *n* indicates the number of neurons. **(C)** Temporal profiles of local and global responses. Top and bottom rows present example neuron and average population responses, respectively. **(D)** Comparisons of onset and peak latencies across three response groups and visual areas. Onset latencies are significantly different across three cortical areas (F = 12.77, *p* < 0.001, two-way ANOVA) and neuron groups (F = 11.59, *p* < 0.001, two-way ANOVA). Peak latencies, similar significant effects across three cortical areas (F = 16.13, *p* < 0.001, two-way ANOVA) and neuron groups (F = 7.55, *p* < 0.001, two-way ANOVA). Error bars represent SEM. Response latencies of high-and low-SF-preferring neurons to grating stimuli are also shown for reference. **(E)** Latency difference between low-and high-SF preferring neurons (>3 cycles/° − <1 cycle/°). **(F)** Latency differences among high-SF global, low-SF global and high-SF local responses. Low-SF global remained the shortest latencies in its onset and peak responses. The direct comparison of latency differences among the three neuronal response groups further demonstrated that the prolonged onset latencies were the primary factor temporally separating processing among the three groups.

We plotted the temporal profiles of these local and global neuronal responses of global–local stimuli, and compared their onset and peak latencies both within and across V1, V2, and V4 (**Figure 4CD and Figure S6**). Onset latency was defined as the time at which the response exceeded the baseline by three standard deviations^6^. We found that, within each cortical area, low-SF coarse global responses occurred earliest (mean ± SEM: 45.6 ± 1.2, 49.1 ± 1.3 and 51.0 ± 1.2 ms in V1, V2 and V4, respectively), followed by fine-local responses (48.6 ± 0.8, 51.4 ± 1.1, 53.5 ± 0.8 ms), whereas high-SF global responses surprisingly exhibited the longest latencies (53.7 ± 1.6, 56.0 ± 2.1, 57.1 ± 1.5 ms) (two-way ANOVA, F = 11.59, *p <* 0.001). As expected, these distinct response latencies across the three response categories increased monotonically from V1 to V4 along the visual hierarchy (two-way ANOVA, F = 12.77, *p <* 0.001). We also examined peak response latencies and found high-SF global responses again exhibited the longest latencies (68.5 ± 2.3, 70.2 ± 2.6 and 72.4 ± 1.4 ms in V1, V2 and V4, respectively), compared with high-SF local (63.1 ± 0.8, 65.1 ± 1.2, 69.3 ± 1.1 ms) and low-SF global (56.7 ± 1.5, 63.6 ± 1.2, 67.3 ± 1.5 ms) within each cortical area (two-way ANOVA, F = 7.55, *p <* 0.001) (**Figure 4D**). Note that the temporal separation of global and local processing was greatest in V1. Furthermore, across cortical areas, neuronal response latencies of high-SF global were also significantly longer than those to high-SF sinusoidal grating stimuli (Onset latency: two-way ANOVA, F = 20.12, *p <* 0.001; post hoc Tukey–Kramer correction, *p =* 0.03. Peak latency: two-way ANOVA, F = 13.83, *p <* 0.001; post hoc Tukey–Kramer correction, *p =* 9.50 × 10⁻^3^). By contrast, no statistical differences between low-SF global and low-SF sinusoidal grating stimuli (post hoc Tukey–Kramer correction, onset latency: *p =* 0.83; peak latency: *p =* 0.78), and between high-SF local and high-SF sinusoidal grating stimuli (post hoc Tukey–Kramer correction, onset latency: *p =* 0.27; peak latency: *p =* 0.93).

It is also important to note that, although low-SF responses exhibited shorter latency than high SFs, the latency difference between low-and high-SF responses to sinusoidal–grating stimuli is much shorter than those between coarse and fine global responses to global–local stimuli (**Figure 4EF**). This difference further support more complex computation in cortical cellular processing of global–local stimuli (**Figure 3F**). Furthermore, except V1 low-SF global, response duration between onset and peak latencies is close to each other among the three visual cortices, indicating the processing time from onset to the peak responses essentially very similar at each cortex. Thus, differences in onset response latency separated the three groups in their responses to global–local stimuli within each cortical area (**Figure 4F**). Together, response latencies in parafoveal V1, V2 and V4 revealed closely spaced but distinct processing stages: coarse global information first, fine local information next, and fine global information last. This processing sequence emerged as early as V1 and persisted through higher cortical areas V2 and V4, with latencies increasing progressively along the visual hierarchy.

### Foveal two-photon imaging across V1, V2, V4, and IT

Primate vision is predominantly foveal, yet the cortical mechanisms underlying global and local perception in the fovea remain unclear (**Figure 1 and Figure S1**). Using two-photon calcium imaging, we recorded a large population of orientation-selective neurons in the foveal regions of V1, V2, and V4 at eccentricities between 1° and 2° from four awake macaques (1,045 neurons in V1; 902 in V2; 1,111 in V4) (**Figures 1F and 5A**). We also examined IT, a high-level visual area at the end of the ventral visual stream that has been classically proposed to contribute to early coarse global perception^9,11,13,40^. We successfully recorded 47 orientation-selective IT neurons within an object-selective patch identified by fMRI in one awake macaque. Compared with V1, V2, and V4, fewer orientation-selective neurons were observed in this IT patch examined (**Figure 5A**), likely because IT neurons are located in the latter stages of visual processing, having large RFs spanning both foveal, parafoveal regions and beyond, and are preferentially driven by more complex natural stimuli.

**Figure 5.**
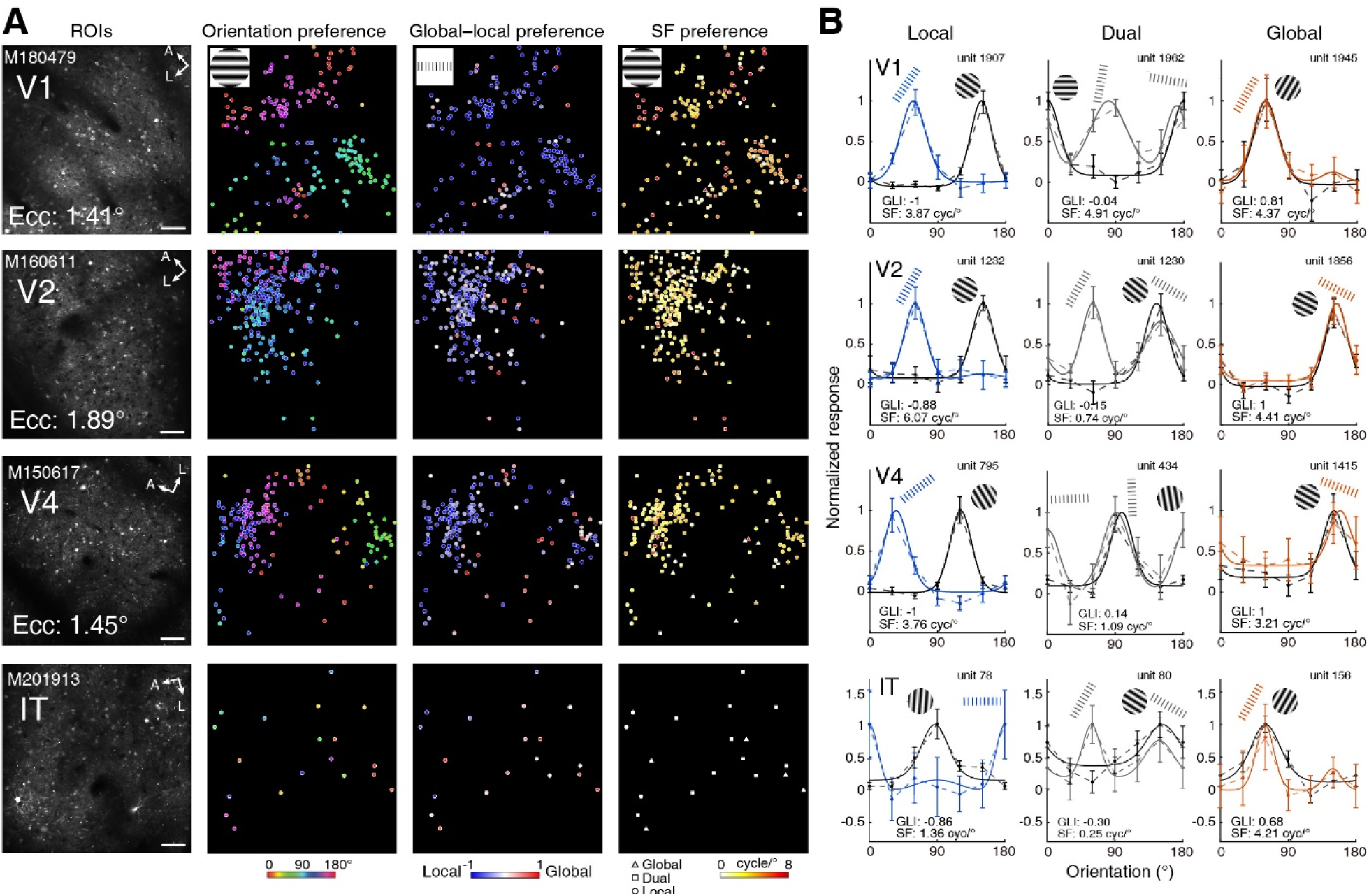
Local-, dual-, and global-selective neurons in foveal regions across V1, V2, V4, and IT revealed by two-photon imaging. **(A)** Two-photon imaging in V1, V2, V4, and IT. The leftmost column shows ROIs in V1, V2, V4, and IT within retinal eccentricities between 1°–2°. L, lateral; A, anterior; Ecc, eccentricity. The right columns show individual cell distributions in grating orientation preference maps, global–local preference maps, and grating SF preference maps. Scale bars, 100 μm. **(B)** Examples of local-, dual-, and global-selective neurons in V1, V2, V4, and IT. Stimulus icons indicate the preferred stimuli for each neuron. GLI, global–local index; SF, neuron preferred SF measured with sinusoidal grating stimuli. See Figure S7A for additional examples of global-selective neurons with low-SF preference.

We observed local-, dual-, and global-selective neurons in the foveal regions as early as V1 and across all four cortical areas up to IT (**Figure 5B and Figure S7A**). In contrast to the parafoveal electrophysiology data, where local-selective neurons dominated V1 and V2 but not V4 where global-selective neurons prevalent (**Figure 3**), the two-photon data instead revealed a continued dominance of local-selective neurons in foveal V4, whereas a dominance of global-selective neurons in IT (**Figure 6A**). Note that the foveal local-selective dominance in V4 is consistent with V4 population responses within 2° eccentricity revealed by intrinsic-signal optical imaging (**Figure 2C**). In addition, local-selective neurons also exhibited significantly stronger calcium signals than global-selective neurons in foveal V1, V2, and V4 (**Figure 6B**). To quantify whether these neurons are organized into distinct functional domains, we calculated their cluster indices across different cortical distances^34,41^ (**Figure 6C**). Because local-selective neurons were both dominant and widely distributed within the foveal imaging windows, their cluster indices remained higher (above 1). In contrast, the cluster indices of global-selective neurons were consistently below 1 across all cortical distances and foveal visual areas, indicating that these global-selective neurons were not spatially clustered but instead were intermingled with local-and dual-selective neurons in a salt-and-pepper pattern.

**Figure 6.**
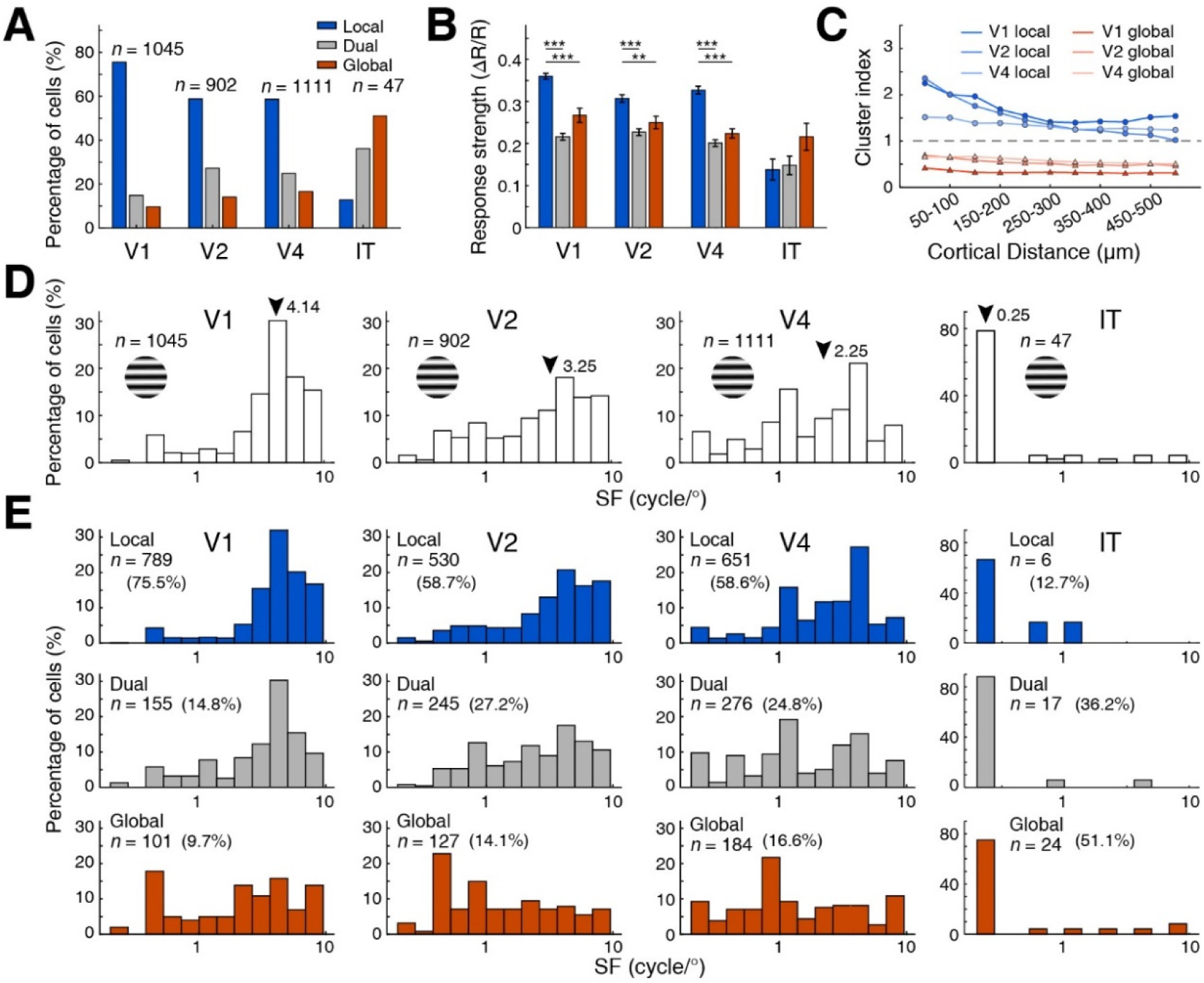
Fine and coarse global encoding together with fine local encoding in foveal regions across V1, V2, V4, and IT. **(A)** Proportions of local-, dual-, and global-selective neurons in foveal regions across V1, V2, V4, and IT. *n* indicates the number of recorded neurons in each area. **(B)** Comparison of response strength among local-, dual-, and global-selective neurons to global–local stimuli across V1, V2, and V4. Two-way ANOVA revealed significant effects across cortical areas (F = 10.67, *p* < 0.001) and neuron types (F = 105.96, *p* < 0.001). Post hoc Tukey–Kramer correction: \*\**p* < 0.01; \*\*\**p* < 0.001. **(C)** Comparison of clustering indices for local-and global-selective neurons in V1, V2, and V4. Clustering indices for global-selective neurons are significantly below 1 at all distances, whereas those for local-selective neurons are above 1 (*p* < 0.001, bootstrap resampling with 1,000 iterations). **(D)** Distributions of preferred SFs for all neurons recorded using two-photon imaging in V1, V2, V4, and IT. Arrow indicates median SFs for each cortex. *n* indicates the number of neurons. **(E)** Distributions of preferred SFs for local-, dual-, and global-selective neurons, respectively, in V1, V2, V4, and IT. *n* indicates the number of neurons.

Consistent with expectations, preferred SFs in the recorded foveal regions also decreased along the visual hierarchy (median SFs: 4.14 cycles/°, 3.25 cycles/°, 2.25 cycles/° across V1, V2 and V4; one-way ANOVA, F = 123.22, *p <* 0.001) (**Figure 6D**), however, they are substantially higher than those recorded in parafoveal regions (**Figure 3B**). Consistent with hierarchical progression and the larger RFs of IT neurons, the preferred SF in IT was lowest (median SF: 0.25 cycles/°) among four recorded visual areas (**Figure 6D**). Notably, among global-selective neurons, we identified a subset of IT neurons (3/24) with high-SF preferences (>3 cycles/°), suggesting that fine global information can be preserved and encoded even at the highest stage of the visual hierarchy. Moreover, in addition to global-selective neurons with high SFs, a subset of global-selective neurons with low SF preference was also observed in the foveal region within each cortical area across cortices (**Figure 6E**). Low-SF-preferring local-selective neurons were also found in the fovea of V1, V2 and V4 (**Figure S7B**). In V1 and V2, these neurons exhibited second but small response peaks to gratings above 3 cycles/°, suggesting possible functional interactions among neuronal populations with different SF preferences.

Together these results indicate that fine spatial information of both local and global features is fully encoded in the fovea as early as V1 and preserved hierarchically up to IT, whereas the coarse spatial information of the global, while present in the fovea from V1, is dominant in higher cortical stages, particularly in IT of the hierarchy.

### Dynamic coherent integration of global and local information

By examining neuronal responses in foveal and parafoveal regions of four successive cortices from V1 to IT, we found that both fine and coarse global information are jointly represented with fine local information as early as V1 and within each following cortical area (**Figure 7A**). However, at the neuronal level, local-selective neurons dominated V1, and their prevalence declined more heavily across the cortical hierarchy than across retinal eccentricity, whereas the proportion of low-SF global-selective neurons increased markedly (**Figure 7AB**). This pattern is consistent with the rapid hierarchical expansion of RF size and the corresponding decline in visual acuity along the visual hierarchy. For global information, the proportion of fine global representation was at least twice that of coarse global in V1, whereas the opposite pattern was observed in higher visual cortical areas (**Figure 7B**). Importantly, the proportion of fine global representation remained relatively constant from V1 through IT, suggesting that fine-global information of V1 is preserved in foveal representations in parallel with the marked increase in low-SF-defined coarse-global information in parafoveal representations. Accordingly, the ratio of high-SF to low-SF global-selective neurons decreased markedly from V1 to IT, primarily because of the increased proportion of low-SF global-selective neurons. Thus, global representation spans a broad range of SFs (**Figure 3C and Figure 6E**), rather than being restricted to low-SF bands, supporting the view that global form and local features are represented in parallel, but with different dependencies on SF range, retinal eccentricity and cortical stage.

**Figure 7.**
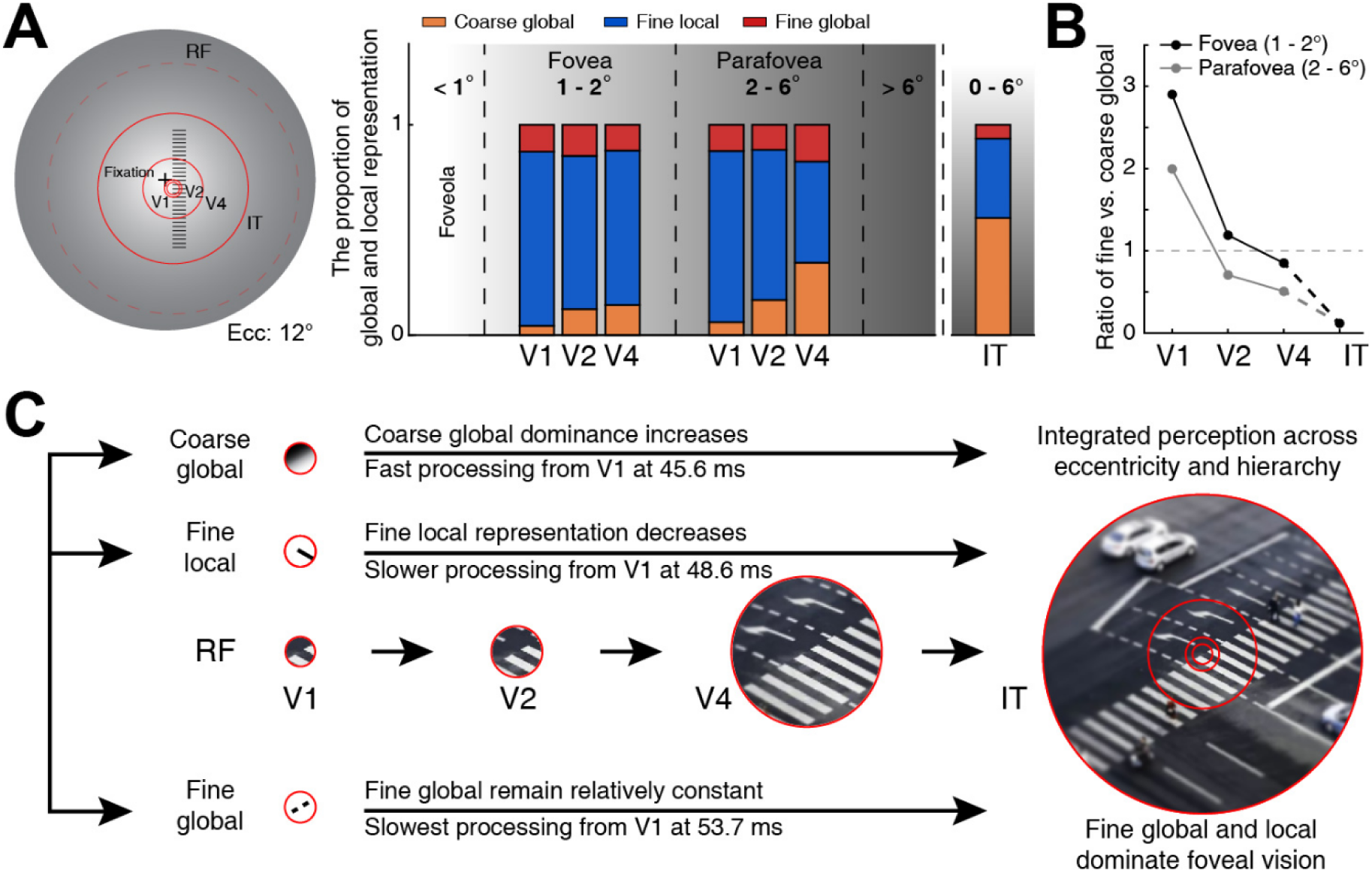
Representations of coarse-global, fine-global and fine-local information across eccentricity along the ventral visual hierarchy. **(A)** Proportion of fine-global, fine-local and coarse-global representations. Left, relative eccentricities and sizes of stimuli. The large solid red circle indicates IT RF covers 0–6° eccentricity. The dashed red circle illustrates IT RF which often above 20° to nature images^45^. Right, proportion of fine-global, fine-local and coarse-global representations in the fovea and parafovea across cortices from V1 to IT. Note that the proportion of fine global is relatively constant when compared with rapid increase of coarse global across cortices from V1 to IT. **(B)** Ratios of fine global to coarse global in fovea and parafovea along visual hierarchy. **(C)** A framework of integrated visual perception of global and local information through different RFs across eccentricities along the visual hierarchy. With a street zebra crossing picture as an example, schematic illustrating cortical neuronal processing of global and local features in a sequence of a coarse-global → fine-local → fine-global information in the foveal and parafoveal regions at each cortex from V1 to IT. RFs sizes were schematized based on Kravitz et al.^28^.

Together, our recordings across V1, V2, V4, and IT at 1°–6° eccentricities show that fine-and coarse-global forms and fine-local parts are represented in parallel at each cortical area from V1 to IT. Within fovea 2° eccentricity high-SF defined fine global and local information is relatively preserved along the hierarchy. At each cortical area, processing unfolded sequentially from coarse-global to fine-local to fine-global representations, within a tightly defined timing sequence, supporting a dynamic neuronal framework of integrated vision across both retinal eccentricity and visual hierarchy (**Figure 7C**). These neuronal results are consistent with the prediction from SF power analysis across different RF sizes of the visual hierarchy (**Figure S1C**). Specifically, the sharpness of both global form and local detail is encoded predominantly by high-SF neurons, especially in foveal regions across the visual hierarchy, supporting stable but slow high-resolution central vision^2,3,42,43^. In contrast, the rapidly increasing of low-SF-preferring neurons representing coarse global form across eccentricity and hierarchy, are consistent with blurred but fast perifoveal and peripheral vision over larger visual fields^23,24,39,44^.

## Discussion

In our daily life, we can effortlessly resolve both fine global shapes or forms and fine local details of small objects in central vision (including foveola, fovea and parafovea), while also grasping the gist of multiple items in the surrounding scene (perifovea and periphery). However, a century-old question, dating back to the Gestalt psychologists and their proposal of “global precedence”^4,5^, remains unresolved: how are global form and local features represented at the neuronal level to generate coherent visual perception across the visual field? Current hierarchical theories of visual feature integration, including local-to-global and coarse-to-fine frameworks^9–11,13,14,25–28^, have not specifically explained how fine global forms and local details are jointly represented by cortical neurons in primate foveal vision. Our findings extend previous work on parafoveal and peripheral vision and establish a new cortical framework for understanding small-object perception in central vision.

### Multi-scale global and local processing across eccentricity and cortical hierarchy

The Gestalt notion of “global precedence”^4,5^ gave rise to classical models of visual processing in which global form perception is associated with low-SF information and local feature analysis with high-SF components^6,8,9,46^. This view has shaped dominant frameworks in which coarse global structure is extracted early and subsequently refined by local analysis, for example through “hierarchy and reverse hierarchy” loops in which IT represents the global whole before fine local details are recovered from V1 inputs^9,11^. However, both natural image statistics^20,21^ (**Figure 1B**) and the global–local stimuli used here (**Figure S1A**) indicate that global structure is not confined to low SFs, but instead spans a broad range of spatial scales, including high-SF components. Consistent with this, our neuronal recordings from foveal and parafoveal regions showed that high-SF-preferring neurons were not limited to their expected role in local processing; a substantial high-SF population also encoded fine global information across ventral cortical areas. In foveal vision, neurons with small RFs and high-SF sensitivity support not only fine local analysis but also the encoding of fine-scale global form, enabling global and local information to coexist in high-resolution representations. Thus, these findings revise conventional coarse-to-fine accounts by showing that in addition to coarse spatial processing, global perception also engages fine-scale processing in the foveal vision. They provide a neuronal basis for the capacity of central vision to resolve both precise global form and fine local detail from as early as V1 through IT, supporting high-acuity visual recognition in central vision (**Figure 1A**).

Current feedforward processing theories proposed that visual information advances from simple to increasingly complex features and culminates in higher cortical areas associated with conscious perception. Our results refine these classical hierarchical frameworks, which have emphasized a progressive transition from local representations in early visual cortex to global and complex representations in higher visual areas^11,25,26,28,29^. Although these feature integration and assembly theories suggest that object recognition may culminate in IT and other higher cognitive areas, we found that basic two-dimensional global and local spatial information is already encoded in parallel as early as V1 and across all stages of the ventral visual pathway, with a gradual shift towards global dominance along the hierarchy (**Figure 7A**). This dynamic multi-scale encoding, together with hierarchical enlargement of RFs, may allow the visual system to sample over increasing regions of visual space while preserving stable high-resolution perception of global forms and local details in the fovea (**Figure 1A**). In other words, IT neurons, which exhibit stimulus-dependent effective RF sizes^45,47^, may encode fine global form together with fine local detail within the foveal portion of their RFs, whereas neurons with larger RFs may represent coarse global form across the entire visual field. Beyond the expected high-SF local-selective neurons, the presence of both fine-and coarse-scale global-selective neurons are already present in V1, consistent with previous observations that V1 neurons can encode natural complex structures, synthetic images and internal sharp illusory shapes^33,36,48–50^. Previous studies have also shown that IT neurons encode fine-grained information within facial stimuli by changing SF, stimulus size, or facial expressions^32,40,51^, consistent with both our present findings in both foveal and parafoveal regions and our previous evidence that fine-scale information is preserved in the parafoveal regions through successive stages of the ventral visual pathway^6,37^. Recent high-resolution fMRI further indicates that multiple visual areas, including IT and other higher-order regions, can be robustly driven by stimuli presented within the foveola and fovea^52^. At eccentricities within 1°, IT cells exhibit significant selective responses to small geometry forms^53,54^, consistent with the observation that a subset of IT cells have small RFs^47,53^. However, these studies did not explicitly dissect global and local features or distinguish their associations with fine-and coarse-scale representations. Our findings extend these important observations by demonstrating that IT neurons encode high-SF-defined global form in addition to coarse global and fine local features. Importantly, the presence of fine-scale global processing as early as V1 suggests an early source for high-SF global representations in subsequent cortical stages up to IT. Notably, whereas the enormous RFs of IT neurons place foveal regions in their RF centers, V1, V2, and V4 are retinotopically organized, each containing distinct high-resolution cortical representations of the fovea.

### Closely spaced temporal processing of global form and local features

In primate retina, the fovea cones exhibited slower light responses up to 30 milliseconds than peripheral cones^2,43^, supporting high acuity but slow visual perception^42,44^. Regardless, our neuronal latency results from parafovea recordings accord with classical accounts of coarse-to-fine perception, which propose that rapid processing of low-SF global information provides a causal basis for the perceptual precedence of global form^9,11,14^. We refined this framework by distinguishing fine global processing from coarse global processing: high-SF global-selective neurons exhibited the longest response latencies across parafoveal representations (2°–6° eccentricity) in V1, V2 and V4. The latencies of low-SF global, high-SF local, and high-SF global responses all increased monotonically along the visual hierarchy, consistent with feedforward propagation and progressive integration across successive cortical stages^28–30^. However, these response latencies were all below ∼100ms, the timescale required for object identification and recognition^9,55,56^. The response delays among these groups were tightly clustered within only a few milliseconds (**Figure 4 and Figure S6**), and how such small temporal differences contribute to rapid categorization and identification of small objects remains unclear. Regardless, these distinct yet closely spaced responses to global form and local detail suggest that meaningful visual representations emerge early within the ventral visual hierarchy prior to extensive engagement of higher cognitive cortices^9,12^. Furthermore, our finding that the temporal processing of fine and coarse global forms by high-and low-SF-preferring neurons emerges as early as V1 and persists across successive cortical stages does not support reverse-hierarchy models in which coarse-to-fine processing is proposed to proceed from IT to V1^9,11^. Although the precise role of top-down feedback from IT and other higher cortical areas to earlier visual cortices remain unclear, it may contribute to higher-level functions such as attentional modulation and size-and position-invariant object representation^54,57,58^. Importantly, SF tuning to drifting gratings is itself dynamically time-dependent^59,60^, raising the possibility that coarse-global, fine-local and fine-global representations may likewise evolve over time. Consistent with this possibility, recent evidence suggests that early IT responses encode relatively simple features, whereas later responses encode more complex features^61^. Psychophysical studies have shown that removing low-SF information diminishes the temporal advantage of global processing^62,63^, consistent with our observation that low-SF global neuronal responses occurred before high-SF fine-local and fine-global responses. Together, these parafoveal response latency findings reveal a tightly spaced temporal sequence of cortical processing, unfolding within each area from coarse-global to fine-local to fine-global representations, and across cortical stages from V1 through IT.

### The advantage of fine global representation for small objects in foveal V1

V1 neurons are not merely simple orientation and spatial-frequency filters^33,48,64^, as suggested by responses to sinusoidal grating stimuli, but also have the capacity to integrate local information into precise global representations of small objects. The simplest example of visual feature construction is the de novo generation of orientation and direction selectivity through spatiotemporal integration of collinearly organized inputs from the lateral geniculate nucleus by V1 layer 4 neurons with simple RFs^65–69^. Many V1 neurons also exhibit complex RF properties that encode line ends, angles and other local configurations^25,33,48^. Furthermore, V1 neurons can signal figure–ground segregation^70,71^.

Recent findings further demonstrate that this integrative capability critically depends on cross-laminar interactions across V1 sublayers. For example, cascaded normalization mechanisms across V1 sublayers substantially transform spatial integration from input to output layers^72^, whereas high-SF information in primate V1 is not merely inherited from feedforward P-pathway inputs, but can be selectively amplified by local recurrent circuits in superficial output layers^73^. More recently, studies using natural images and complex visual stimuli have shown that V1 neurons respond robustly to richer visual structures than simple Gaussian–Gabor filter models would predict^33,36,48–50,74^. Together, these findings indicate that V1 provides a more flexible substrate for spatial integration than classical filter-based descriptions alone would suggest.

This computational advantage and capacity for basic global information processing at early cortical stage of the visual hierarchy is especially important for small-object representation in foveal vision. Small objects fall within the high-acuity foveal map, where V1 neurons have small RFs and preserve precise retinotopic relationships. Thus, retinotopic information about fine global form and local detail can be represented in parallel before spatial information is degraded by pooling across eccentricity in higher cortical areas for progressively coarse global representations (**Figure 7**). This view is consistent with our finding that neurons selective for global information over increasing visual space than local line elements across V1, V2, and V4 exhibited more nonlinearity when tested with sinusoidal gratings (**Figure 3F**). Note that large objects extend across foveal, parafoveal and peripheral regions will require progressively broader, coherent integration over larger RFs in higher visual cortices, which must have to draw on distributed inputs from corresponding retinotopic regions of early visual areas including V1. Such integration is essential for representing large-scale object structure, but it comes at the cost of reduced spatial resolution in the parafoveal and peripheral regions. From this perspective, higher visual areas are well suited for integrating large objects and supporting category-level representations, whereas foveal V1 provides a uniquely precise substrate for jointly resolving the global form and fine local details of small objects with high spatial precision.

### Several limitations of the study

First, selective attention can bias perception toward either global structure or local detail^75,76^. Because our recordings were performed during passive fixation in awake monkeys, without task demands that selectively emphasized local or global information, we could not determine how attention modulates global–local processing. Thus, the present study addresses feedforward and intrinsic global–local representations rather than selective visual attention. Second, because we used two-dimensional global–local stimuli rather than natural three-dimensional (3-D) objects, our findings should be interpreted as evidence that V1 responses most likely represent two-dimensional global contours/outlines, including fine-scale high-SF global structure such as aligned line endpoints in the stimuli, rather than as evidence that 3-D global objects are represented in V1. In fact, 3-D objects are represented in IT through hierarchical cortical integration^28^. Third, the cortical mechanisms underlying global and local processing in the foveola, corresponding approximately to the central 0–1° of visual space, remain unresolved. This limitation reflects not only the technical challenges of neuronal recording or imaging in the primate visual system^52,77^, but also a more fundamental constraint: the spatial resolution of the foveola exceeds that of standard visual display systems. Consequently, stimuli with sufficiently well-defined global and local features cannot yet be presented at a scale matched to the extremely small RFs of foveolar neurons. Although our recordings did not directly sample the foveola, our near-foveolar dominance of high-SF representations of both global form and local features suggests that fine-scale global structure may extend toward the center of gaze, where the fine global form and local detail of very small objects will be resolved with very high visual acuity, albeit with longer processing times as early as the fovea of primate retina^2,43^.

### A neuronal framework for foveal vision of global and local information processing

Together, these population-and single-neuron results reveal distinctive encodings of global information across multiple SF scales, alongside high-SF local information across retinal eccentricities from V1 to IT. Extending the Gestalt notions of global precedence and filling-in theory^4,5^, our findings show that global processing begins with rapid coarse representations, but also encompasses fine-scale global representations that emerge through coordinated multi-scale and temporally structured processing as early as V1. Although the relationship between neuronal activity and perceptual experience remains difficult to resolve, our findings provide a neural basis for the coherent encoding of fine global form and fine local detail, together with rapid processing of coarse global information in foveal and parafoveal regions within each cortical stage. These results support a dynamic neuronal framework of integrated vision, in which global and local information are jointly constructed across retinal eccentricities particularly in the fovea and along the ventral visual stream. Within this framework, IT neurons, with their large RFs and convergent foveal and parafoveal inputs, may dynamically integrate coarse and fine information to support the categorization and recognition of the same or different objects across a wide range of sizes. For example, coarse global structure may emerge first to support rapid categorization, followed by the progressive hierarchical filling-in of various local details and features that support object identity across spatial scales^40^. More broadly, this framework helps explain how the primate visual system integrates precise central vision with broader peripheral representations to construct coherent perception across large visual field. Together, these findings provide a neural framework for addressing and refining a century-old question in Gestalt theory: how coherent wholes and their constituent parts are jointly constructed in active foveal vision to support fine-scale perception during reading, object recognition and other visually demanding behaviors.

## Acknowledgments

This work was supported by the following grants: Brain Science and Brain-like Intelligence Technology --National Science and Technology Major Project 2022ZD0204600 (to W.W. and S.M.T), the CAS Project for Young Scientists in Basic Research (Grant No. YSBR-113) (to Y.L.L), and the Lingang Laboratory (grant no. LGL-5925-09 to W.W.).

## Author contributions

J.P.Y, Z.Y.C, W.H.X., Y.Z.J.L., Y.L.L., S.M.T performed experiments, and all listed authors contributed to the experiments and data analysis. W.W., Y.L.L, W.H.X., and Y.Z.J.L. wrote the main paper. W.W. designed and supervised the research.

## Materials and Methods

### Psychophysics on human

Psychophysical experiments were conducted in 8 participants (5 males and 3 females; 23–38 years old) with normal or corrected-to-normal vision (**Figure 1E**). All participants provided written informed consent prior to participation, and all procedures were approved by the Ethics Committee of the Institute of Neuroscience, Chinese Academy of Sciences (No. CEBSIT-2023032).

Visual stimuli were presented on a gamma-corrected monitor (Display++, Cambridge Research Systems; 1920 × 1080 pixels, 120 Hz) positioned 67 cm from the eyes, subtending 60.8° × 34.2° of visual angle. Luminance calibration was performed using a SpectroCAL MKII spectroradiometer (Cambridge Research Systems Ltd., Rochester, Kent, UK). Stimuli were generated using Psychtoolbox-3 in MATLAB (MathWorks, Natick, MA, USA). Eye position of the right eye was monitored using an EyeLink 1000 system (SR Research Ltd., Ottawa, Ontario, Canada) at a sampling rate of 1000 Hz, and fixation was required within a 0.5° window.

Global–local stimuli were conceptually similar to the seminal hierarchical stimuli introduced by Navon^7^ and consisted of a global band constructed from short white line elements on a black background. Stimuli were presented within an edge-blurred circular aperture (radius, 1.2°), either at fixation (0°) or along the right horizontal meridian at eccentricities of 2°, 4°, 8°, or 12°. The SFs of the global and local components were fixed at 0.5 and 3 cycles/°, respectively. The global orientation was either near-horizontal (nominal orientations: 3° or 177°) or near-vertical (nominal orientations: 87° or 93°), with a random trial-by-trial jitter of ±0.5°. Local line orientations were selected from the same set of possible orientations, with the constraint that they were near-orthogonal to the global orientation. A two-alternative forced-choice (2AFC) task was used. Each trial began with a text cue (“global” or “local”) instructing participants to report the orientation of the corresponding stimulus component. After a 500 ms fixation period, the stimulus was presented for 200 ms. Participants were instructed to respond as quickly and accurately as possible after stimulus onset by reporting whether the cued orientation was tilted left or right relative to the vertical axis. Reaction time was measured from stimulus onset. Trials without a response within 1000 ms were counted as incorrect and excluded from reaction-time analysis. Each condition was repeated 50 times.

### Intrinsic optical imaging in macaque V1, V2 and V4

All primate experimental procedures were approved by the Animal Care and Use Committee of the Institute of Neuroscience, Chinese Academy of Sciences (No. CEBSIT-2022052). All experimental procedures were also in accordance with the National Institutes of Health Guide for the Care and Use of Laboratory Animals. Five adult rhesus macaques (Macaca mulatta; four males and one female; 6–8 years old; 4.5–10.0 kg) were prepared for simultaneous intrinsic optical imaging of V1, V2, and V4, as previously described^6,34,35,78^.

Visual stimuli were presented on a gamma-corrected CRT monitor (Sony G520; 1280 × 960 pixels, 100 Hz) positioned 57 cm from the eyes, subtending 40° × 30° of visual angle. The fovea and retinal landmarks were projected onto the screen center using a reversing ophthalmoscope^6,79^. Stimuli were generated using Psychtoolbox-3 in MATLAB. Global–local stimuli were constructed from local line elements arranged to form global bands (**Figure 1D**). Local elements were oriented at 45°, 90° or 135° relative to the orientation of the global bands. The SF of the local elements was fixed at 3 cycles/°, whereas the global SF was 0.5 cycles/°, with one cycle consisting of a 1°-wide band and a 1° gap. A dashed-line configuration with 1°-long local lines and a zigzag configuration with 1.4°-long local lines oriented at 45° and 135° relative to the global orientation were also used. Four global orientations (0°, 45°, 90°, and 135°) were tested. As a control, drifting sinusoidal gratings were presented at four orientations (0°, 45°, 90°, and 135°) and eight SFs (0.25, 0.5, 0.75, 1, 2, 4, 6, 8 cycles/°), with temporal frequency fixed at 1 Hz. Retinotopic and eccentricity maps were obtained using drifting light bars (0.5° width) moving in orthogonal directions at 0.08 Hz (2.33°/s), covering 28° × 28° of visual space, presented monocularly^6^.

Intrinsic signals were computed as the fractional change in reflectance ((R_1_ - R_0_)/R_0_), where baseline (R_0_) was defined as the mean signal during the 1 s pre-stimulus period, and response (R_1_) as the mean signal during 2-6 s after stimulus onset. Differential maps were generated by pixel-wise subtraction of responses to paired stimuli (e.g., 0° - 90° or 45° - 135°). Orientation preference maps were constructed using a vector summation method (**Figure S2A**). For visualization, maps were high-pass filtered (1.1–1.2 mm in diameter) and smoothed (85–323 μm in diameter) using circular averaging filters to reduce noise without distorting signal structure^6,34,35,78^. For quantitative analysis, pixel reliability across trials was quantified as cross-trial variance^6,80^. Based on the histogram of the variance map, an objective threshold was determined as the intersection of a Gaussian and a power law fit of the distribution. Pixels with large cross-trial variance higher than the threshold were overlaid by a gray mask, and were not used in further quantitative analysis. Retinotopic maps were derived using Fourier analysis of responses to moving-bar stimuli^6,81^. The response phase at the stimulus frequency was extracted for each pixel, and hemodynamic delays were corrected by subtracting phase maps obtained from opposite motion directions. Phase values were converted into azimuth and elevation maps (-14° to 14° visual angle), from which eccentricity maps were computed. Response profile analysis was performed to extract orientation components from differential maps^6,35,82,83^ (**Figure S2B**). Global–local preference maps were generated by extracting the peak orientation of the response profile at each pixel, with averaging within an 800 μm neighborhood in response to global–local stimuli (**Figure 2BC**).

### Electrophysiology in macaque V1, V2 and V4

All primate experimental procedures were approved by the Animal Care and Use Committee of the Institute of Neuroscience, Chinese Academy of Sciences (No. CEBSIT-2022052). All experimental procedures were also in accordance with the National Institutes of Health Guide for the Care and Use of Laboratory Animals. Three adult rhesus macaques (Macaca mulatta, all males, 6–10 years old, 6.9–12.0 kg) were trained for awake electrophysiology, as previously described^6,34^. Head fixation was achieved using a custom biocompatible titanium headpost, and a custom designed biocompatible PEEK recording chamber was implanted over visual areas V1, V2, and V4. A grid system (0.8 mm spacing) was used to guide electrode penetrations and map recording locations, allowing estimation of retinal eccentricity across penetrations. The dura was replaced with an artificial dura to facilitate stable and perpendicular electrode access.

Visual stimuli were generated using Psychtoolbox-3 in MATLAB and presented on a gamma-corrected LCD display (Display++, Cambridge Research Systems), positioned 114 cm from the eyes. Eye position was monitored using an infrared eye-tracking system (Eyelink 1000, SR Research) at 500 Hz, and fixation was required within a 1° window. Each trial began with 800 ms fixation, followed by stimulus presentation (1000–1500 ms), and an additional 500 ms fixation period after stimulus offset. Stimuli were presented either full-field or within the neuron’s RF. Drifting sinusoidal gratings were presented at 12 orientations (0°–165° in 15° steps, moving perpendicularly to their orientations) to determine orientation preference. Gratings with a range of SFs (0.125, 0.25, 0.5, 1, 2, 3, 4, 6, 8 cycles/°) were presented at the neuron’s preferred orientation to determine SF preference. Temporal frequency was set at 1–5 cycles/s. Global–local stimuli were also presented at the same 12 orientations, with these orientations referring to the global component (SF: 0.5 cycles/°). The local component (SF: 3 cycles/°) was always oriented 90° relative to the global component.

Neuronal activity was recorded using either single tungsten electrodes or 16-channel linear array electrodes (U-probes, Plexon Inc.; 150 μm inter-contact spacing), advanced with a microdrive (Narishige). Signals were acquired using Omniplex recording system (Plexon Inc.) at a sampling rate of 40 kHz. Spike detection and sorting were performed offline using standard methods published previously^6,34,84^. Data analysis was performed using custom MATLAB scripts. Neuronal response to a given stimulus is the averaging spike firing rate (0 ms to 1000 ms from stimulus onset). Spike density functions (SDF) were obtained by convolving spike trains with a Gaussian kernel (σ = 5 ms)^85^. The response threshold was defined as the mean spontaneous firing rate (−200 ms to 0 ms before stimulus onset) plus three times its standard deviation. Neurons whose SDFs did not exceed the threshold within 100 ms after stimulus onset were excluded from latency analysis. Onset latency was defined as the first time point at which the SDF exceeded the threshold for at least 10 ms, whereas peak latency was defined as the time point of the maximum SDF response within 200 ms after stimulus onset for responses that exceeded the predefined threshold. Because responses of a few neurons were not sufficiently stable or consistent to meet these criteria, the numbers of neurons included in the onset-and peak-latency analyses could differ.

Orientation tuning curves were obtained from responses to grating stimuli. The orientation selectivity index (OSI) was calculated by a vector summation method^86^:

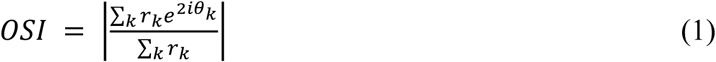

where *θ*_k_ and *r*_k_ are the stimuli orientation and the average firing rate, respectively. Neurons with OSI < 0.1 were excluded from further analysis. Responses to grating stimuli were fitted with a von Mises function to determine each neuron’s preferred orientation. The von Mises function is defined as:

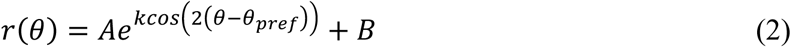

where *A* scales the height of the tuning curve, *k* determines the tuning bandwidth, *θ* is the stimulus orientation, *θ*_pref_ is the preferred orientation, and *B* is the baseline^87^. Fits with R^2^ > 0.5 were considered acceptable.

Responses to global–local stimuli were fitted using a dual-peaked von Mises function, with the two peaks constrained to be 90° apart. The function is defined as:

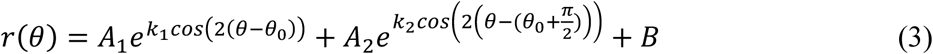

where *θ*_0_ and *θ*_0_ + *π*/2 denote the orientations of the two fitted peaks. *A*_1_ and *A*_z_ are amplitude parameters, *k*_1_ and *k*_z_ control the tuning bandwidths, and *B* represents the baseline. Fits with R^2^ >

0.5 were considered acceptable. Units were excluded from further analysis if the orientations of both fitted peaks deviated by more than 22.5° from *θ*_pref_. For each unit, the fitted response at whichever of *θ*_0_ or *θ*_0_ + *π*/2 was closer to *θ*_pref_ was defined as the response to the global orientation (*r*_global_) of the global–local stimulus, whereas the response at the other peak was defined as the response to the local orientation (*r*_local_). To measure the global and local selectivity of neurons, a global–local index (GLI) was calculated, which is defined as:

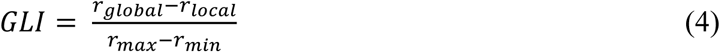

*r*_max_ and *r*_min_ represent the maximum and minimum of the fitted curve. The GLI ranges from −1 to 1, with positive values indicating a preference for global orientation and negative values indicating a preference for local orientation. Neurons were classified as global-selective (GLI > 0.5), local-selective (GLI <-0.5), or dual-selective (intermediate values).

SF tuning curves were fitted with a log-Gaussian function:

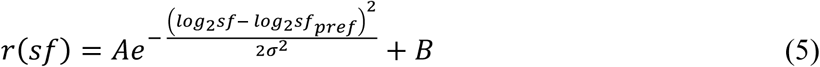

where *A* represents the amplitude, *sf*_pref_ is the preferred SF, *σ* determines the tuning bandwidth, and *B* represents the baseline. Fits with R^2^ > 0.5 were considered acceptable.

RF sizes were measured using methods described in our previous studies^6,84^. For V1 and V2 neurons, RFs were estimated using a white-noise reverse-correlation method with a 16×16 grid of white squares. RF size was defined as the equivalent diameter of the area enclosed by the full-width-at-half-maximum contour of a two-dimensional Gaussian fit to the response map. For V4 neurons, RFs were mapped using 1° grating patches with the neuron’s preferred orientation, and hand-mapped RF size was quantified as the square root of the RF area.

To quantify response linearity, a linearity index was calculated as the ratio of the first harmonic amplitude to the mean firing rate (F1/F0) from SDF of drifting grating stimuli at different SF^84,88^. The max F1/F0 of 3 most preferred SFs was chosen. Higher F1/F0 values indicate more linear responses, whereas lower values reflect more nonlinear responses.

### Two-photon calcium imaging in macaque V1, V2, V4, and IT

All primate experimental procedures followed the guidelines provided by the Institutional Animal Care and Use Committee (IACUC) of Peking University Laboratory Animal Center and were approved by the Peking University Animal Care and Use Committee (No. LSC-TangSM-3). Five adult rhesus macaques (Macaca mulatta; all males; 4-5 years old; 4-9 kg) were used for two-photon calcium imaging, as previously described^33,34,89,90^. Briefly, animals were anesthetized and craniotomies were performed over V1/V2 (2 macaques), V4 (2 macaques), or IT (1 macaque). The dura was opened, and an AAV vector (AAV1.hSynap.GCaMP5g.WPRE.SV40, Penn Vector Core or pAAV.Syn.GCaMP6s.WPRE.SV40, Addgene) was injected at multiple cortical sites. The dura was then sutured, the bone flap replaced and secured, and the skin closed. After 1-2 months, the dura was removed and replaced with a glass cranial window within a titanium recording chamber.

Visual stimuli were generated using a ViSaGe system (Cambridge Research Systems, Cambridge, UK) and presented on an LCD monitor (Acer V173, 80 Hz) positioned 45 cm from the eyes. Monkeys were trained to maintain fixation within a 1° window, monitored using an infrared eye-tracking system (ISCAN ETL-200, ISCAN Inc. USA) at 120 Hz. Each trial consisted of a 1000 ms blank period followed by a 1000 ms stimulus presentation during fixation. Sinusoidal gratings were presented at 6 orientations (0°–150° in 30° steps) and 6 SFs (0.25, 0.5, 1, 2, 4, 8 cycles/°) to determine orientation and SF preference. Global–local stimuli, with a 90° orientation difference between global (0.5 cycles/°) and local (3 cycles/°) components, were presented at the same 6 orientations. For V1, V2 and V4 imaging fields, population RFs were estimated using 0.8° grating patches as previously described^33^, and stimuli were presented within a 10° circular aperture centered on the estimated population RF. In line with recordings from V1, V2 and V4, and given that IT RFs typically encompass central visual field^45,47,91^, for IT recording, stimuli were centered on fixation point within a 10° circular aperture, restricting stimulation to foveal and parafoveal visual field.

Two-photon imaging was performed using a Prairie Ultima IV microscope (Bruker Nano Surfaces Division, FMBU, formerly Prairie Technologies, Middleton, WI, USA) with a Ti:Sapphire laser (Mai Tai eHP, Spectra Physics, Santa Clara, CA, USA). The wavelength of the laser was set at 1000 nm. With a 16× objective (0.8-N.A., Nikon), an area of 850 µm × 850 µm was imaged. Fast resonant scanning (up to 32 frames per second) was used, and images were averaged to yield an effective frame rate of 8 Hz.

Data were analyzed using custom MATLAB scripts as previously described^34^. Imaging frames were first realigned to a template using a normalized cross-correlation-based translation algorithm. ON frames (8 frames after stimulus onset) and OFF frames (8 frames before stimulus onset) were averaged, and differential images were computed as (ON - OFF)/OFF. Neurons were identified using a difference-of-Gaussians filter (5 and 50 pixels), and neuronal responses were quantified as the mean ΔF/F across all pixels corresponding to each neuron. For each neuron, the grating that evoked the maximal response was first identified. Orientation tuning was defined from responses to all orientations at the spatial frequency of this grating. Based on this orientation tuning curve, the OSI was defined as (R_preferred_ - R_orthogonal_)/ R_preferred_^92^, and neurons with OSI < 0.2 were excluded from further analysis. SF tuning was defined from responses to all SFs at the maximal-responding grating’s orientation.

Consistent with electrophysiological analyses, responses to grating stimuli were fitted with a von Mises function to determine preferred orientation, and SF tuning curves were fitted with a log-Gaussian function to determine preferred SF. Responses to global–local stimuli were fitted using a dual-peaked von Mises function, and a GLI was computed to quantify each neuron’s global–local preference. Only neurons for which all three fits yielded R² > 0.5 were included in further analysis. To quantify spatial clustering of functional selectivity, a cluster index (CI) was computed^34,41^:

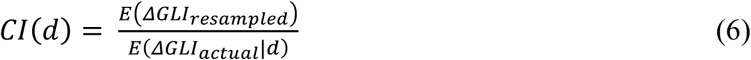

where *d* denotes the cortical distance between neuron pairs, *ΔGLI* is the difference in GLI between two neurons and *E* denotes the expected value. Resampling was repeated 1000 times. CI values greater than 1 indicate that neurons at a given distance are more similar in preference than expected by chance (clustering), whereas values less than 1 indicate relative dispersion.

### Functional MRI in macaque IT

Functional MRI data were acquired on a 3T Siemens Prisma scanner as previously described^93^, using a custom 8-channel surface coil. High-resolution structural images were collected in a separate anesthesia session using a T1-weighted 3D MPRAGE sequence (TR = 2300 ms, TE = 3.8 ms, flip angle = 9°, 224 slices, FOV = 128 × 128 mm, voxel size = 0.5 × 0.5 × 0.5 mm). Functional images were acquired using an echo-planar imaging (EPI) sequence (TR = 2000 ms, TE = 24 ms, flip angle = 80°, 27 slices, FOV = 96 × 96 mm, voxel size = 1.5 × 1.5 × 2 mm). To enhance contrast-to-noise ratio, monocrystalline iron oxide nanoparticles (MION; Molday ION, 0.26 ml/kg, BioPAL) were administered intravenously prior to functional scanning.

Visual stimuli were presented via a video projector (60 Hz, 1024 × 768 resolution) positioned 183 cm from the monkey’s eyes. A block design was used in which each trial consisted of 500 ms stimulus ON followed by 500 ms stimulus OFF, with 24 trials (images) per block. Each run comprised 8 stimulus blocks interleaved with 9 blank blocks (24 s each, fixation only), yielding a total duration of 408 s (204 TRs). For the functional localizer, stimuli from four categories (faces, bodies, objects, and scrambled images; ImageSet, https://github.com/liyipeng-moon/PassiveViewing_in_ML) were presented in separate blocks, with a stimulus diameter of 8°.

Data preprocessing was performed using SPM12 (MATLAB) and included field map correction, motion correction, and spatial normalization. Functional activation was estimated using a general linear model, and category selectivity was assessed using t-contrasts comparing each category against all others.

Functional activation maps were co-registered to high-resolution structural images acquired on the same day and further aligned to the anatomical reference volume using SPM12. Two-photon imaging windows in IT were manually identified in native space based on structural landmarks using 3D Slicer (Slicer Community, Boston, MA, USA). Three-dimensional reconstructions of anatomical and functional data were generated using Blender (Blender Foundation, Amsterdam, the Netherlands).

## Supplementary figures

**Figure S1.**
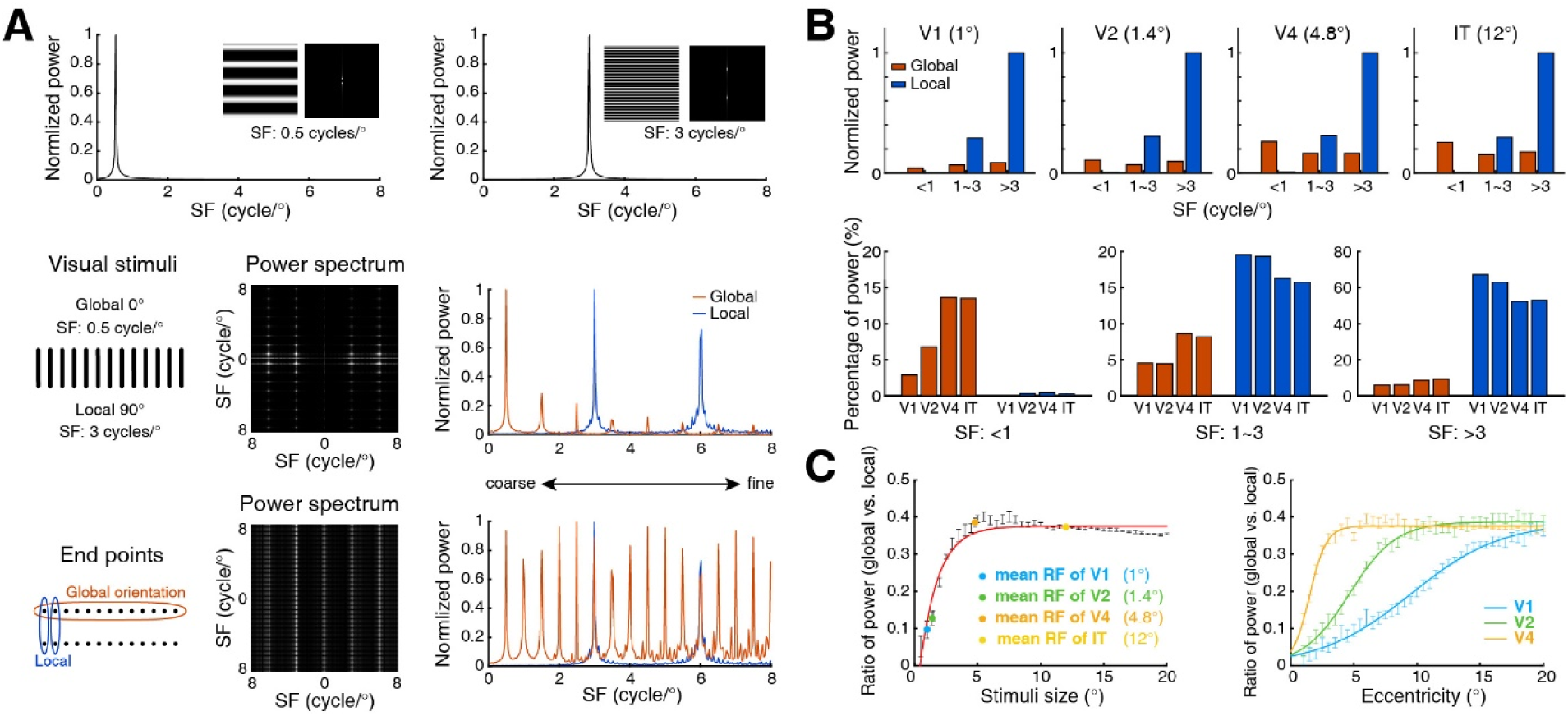
SF power analysis of global and local information through different apertures. **(A)** Spatial power analysis of sinusoidal gratings and global–local stimuli. Sinusoidal gratings are commonly used to measure the SF preferences of visual neurons. As control stimuli, sinusoidal gratings with SFs of 0.5 and 3 cycles/° show single-peak distributions. The SFs of the global and local components of global–local stimuli were set to 0.5 and 3 cycles/°, respectively. The middle panel shows the corresponding spatial power spectrum of a 10×10° global–local stimulus. The right panel shows multiple power peaks across SFs along the orthogonal global and local orientations. These peaks were not restricted to the preset 0.5 cycles/° for the global orientation or the 3 cycles/° for the local orientation, but extended to higher SFs. Spatial power analysis of the line endpoints within the global–local stimuli, shown at the bottom, revealed more peaks across high-SF ranges. **(B)** SF power distributions of the global–local stimuli corresponding to the RFs of V1, V2, V4, and IT neurons. Upper, normalized stimulus power of global and local components for the global–local stimuli in (A) across different SF ranges, with stimulus sizes matched to the mean RF sizes of V1, V2, V4, and IT neurons (1°, 1.4°, 4.8°, and 12°, respectively; derived from Kravitz et al.^28^). Normalized High-SF power dominant in all cortical areas. Lower, the proportion of global and local power between different ranges of SFs across four visual areas. Arrows indicate the distribution trends of SF powers for the global and local, respectively, in corresponding to above RF sizes of V1, V2, V4 and IT neurons. Relative percentage of low-SF powers increase along the visual hierarchy, whereas that of high-SF powers decrease. **(C)** Relationship of spatial power distributions between global and local information across V1, V2, V4, and IT. Left, the ratio of global to local spatial power in the global–local stimuli across various stimulus sizes. Colored dots indicate the ratios at stimulus sizes matched to the mean RF sizes of V1, V2, V4, and IT, demonstrating the power of the global increases along with increasing sizes of global–local stimuli (aperture problem). Right, global-to-local spatial power ratios across retinal eccentricity. The increase of global powers is more rapidly across eccentricity in high visual areas than V1, matching larger increases in RF sizes in V2 and V4 than in V1 along eccentricity^27^. Error bars indicate the SEM.

**Figure S2.**
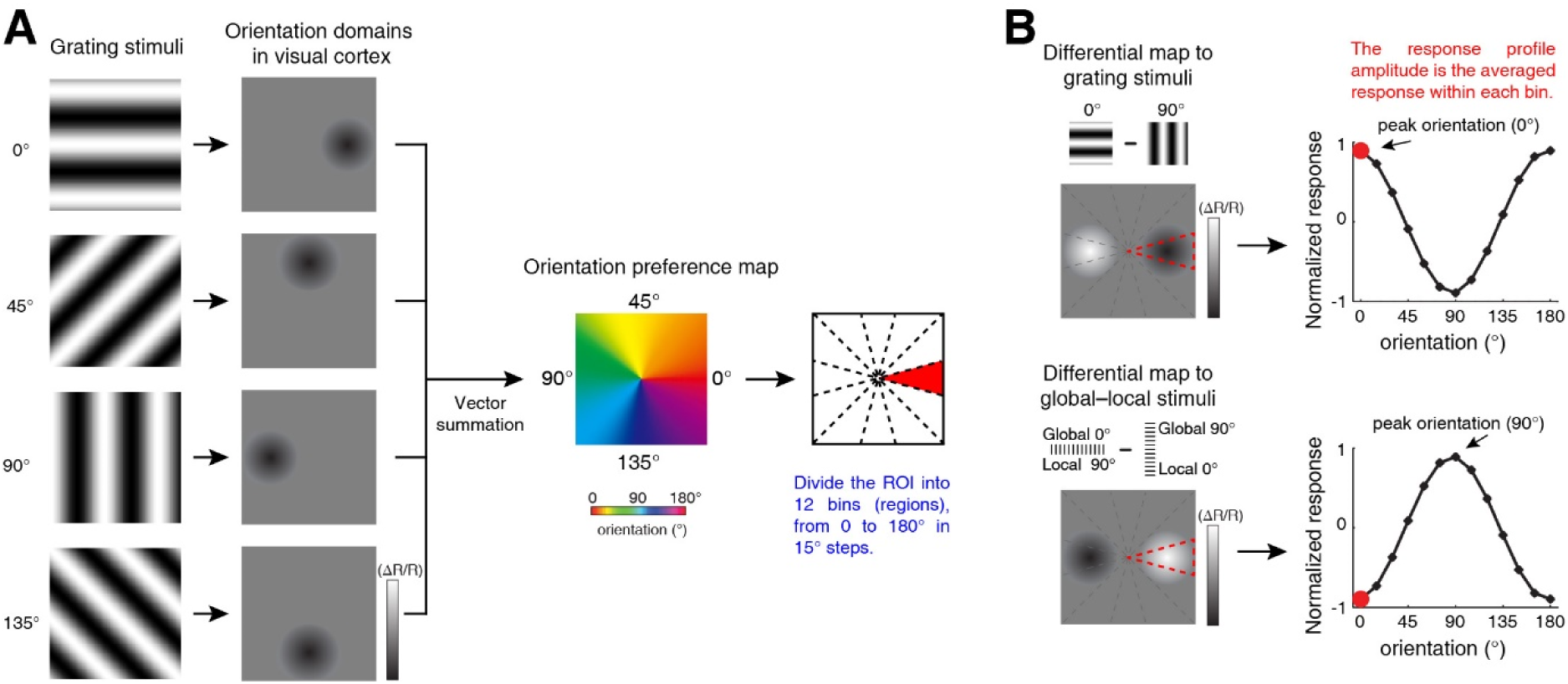
Schematic illustration of the analysis of orientation response profiles for grating and global–local stimuli. **(A)** Responses to grating stimuli at four orientations (0°, 45°, 90°, and 135°) were recorded. Black regions indicate the corresponding orientation domains. An orientation preference map was constructed by vector summation. The imaged ROI was then divided into 12 orientation bins spanning 0°–180° in 15° increments according to the orientation preference map. The region shown in red represents pixels with preferred orientations between −7.5° and 7.5°. **(B)** Differential maps were generated by pixel-wise subtraction of responses to paired grating or global–local stimuli. To obtain the orientation response profile, ΔR/R values were averaged across all pixels within each orientation bin. The red dot in the response profile indicates the mean response amplitude of all pixels within the red region. The arrow indicates the peak orientation of the response profile.

**Figure S3.**
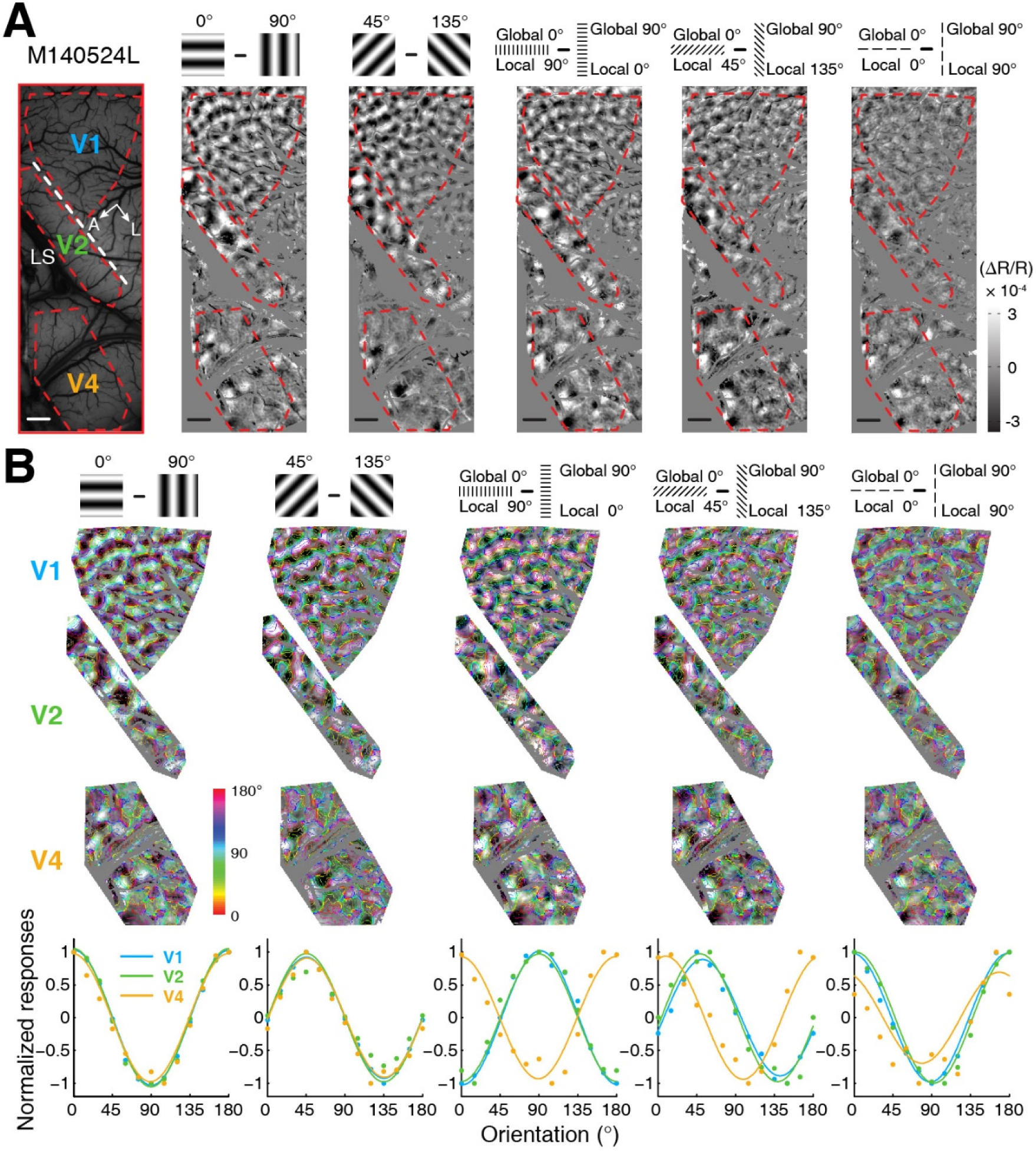
Global and local representations of population responses in parafoveal regions across V1, V2, and V4 in a macaque (M140524). **(A)** Differential orientation maps. Simultaneous recordings from V1, V2, and V4 in response to paired grating stimuli (0° - 90°, 45° - 135°) and global–local stimuli (global: 0° - 90°) with orientation differences between global and local components of 90°, 45°, and 0°. The dashed white line indicates the border between V1 and V2. Red outlines mark the analyzed ROIs in each area. A, anterior; L, lateral; LS, lunate sulcus. Scale bar, 1 mm. **(B)** Orientation response profiles of grating and global–local stimuli. Upper panels show differential maps for paired grating and global–local stimuli overlaid on orientation preference maps. Lower panels show corresponding orientation response profiles for V1, V2, and V4, respectively.

**Figure S4.**
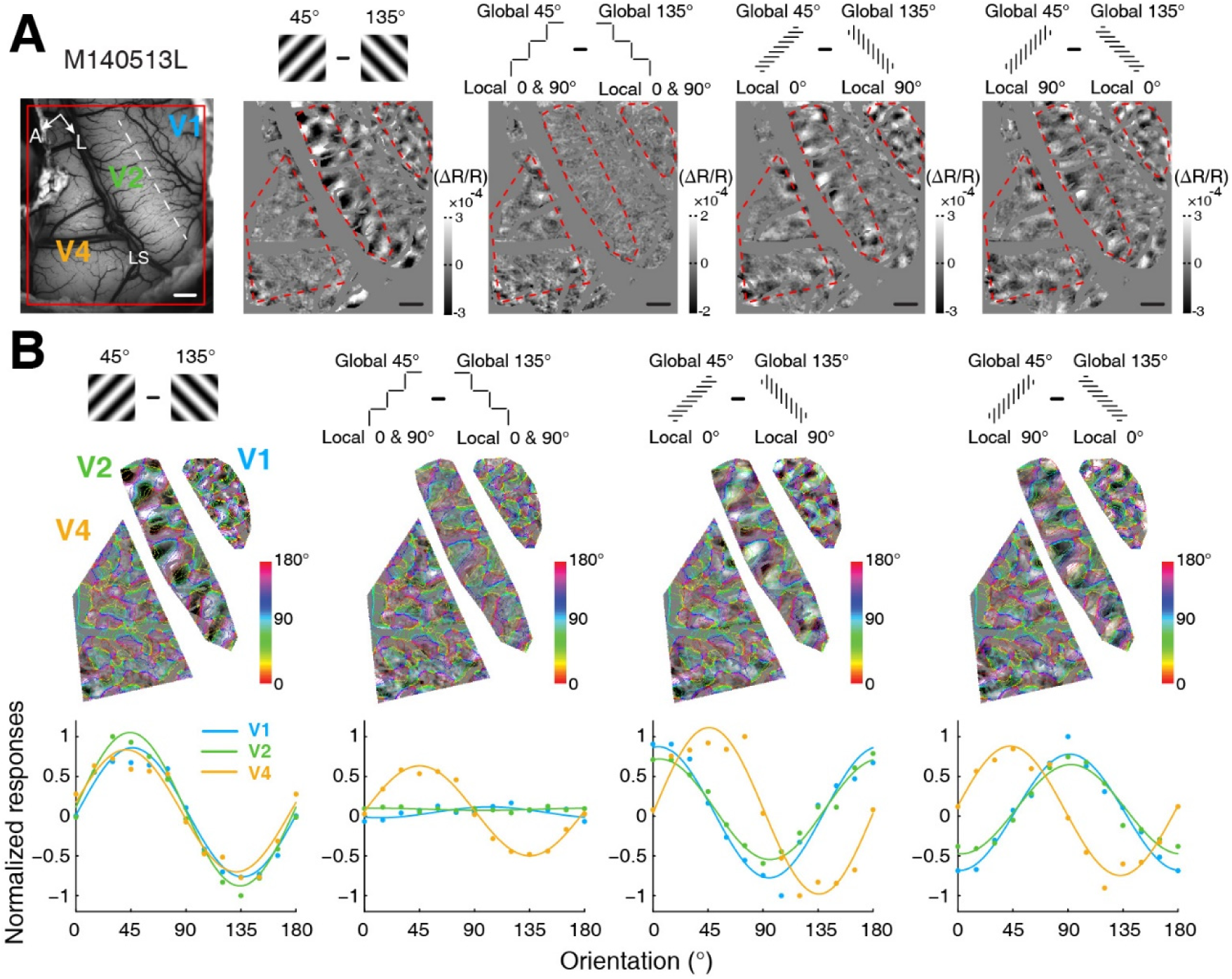
Global and local representations of population responses in parafoveal regions across V1, V2, and V4 in a macaque (M140513). **(A)** Differential orientation maps. The simultaneously recorded V1, V2 and V4 areas in response to paired grating (45° - 135°) and global–local stimuli (global: 45° - 135°) with varies local configurations. The dashed white line indicates the border between V1 and V2. Red outlines indicate the analyzed ROI in each area. A, Anterior; L, lateral; LS, lunate sulcus. Scale bar, 1 mm. **(B)** Orientation response profiles of grating and global–local stimuli. Upper panels, differential maps for paired grating and global–local stimuli superimposed on orientation preference maps. Lower panels, corresponding orientation response profiles in V1, V2 and V4, respectively.

**Figure S5.**
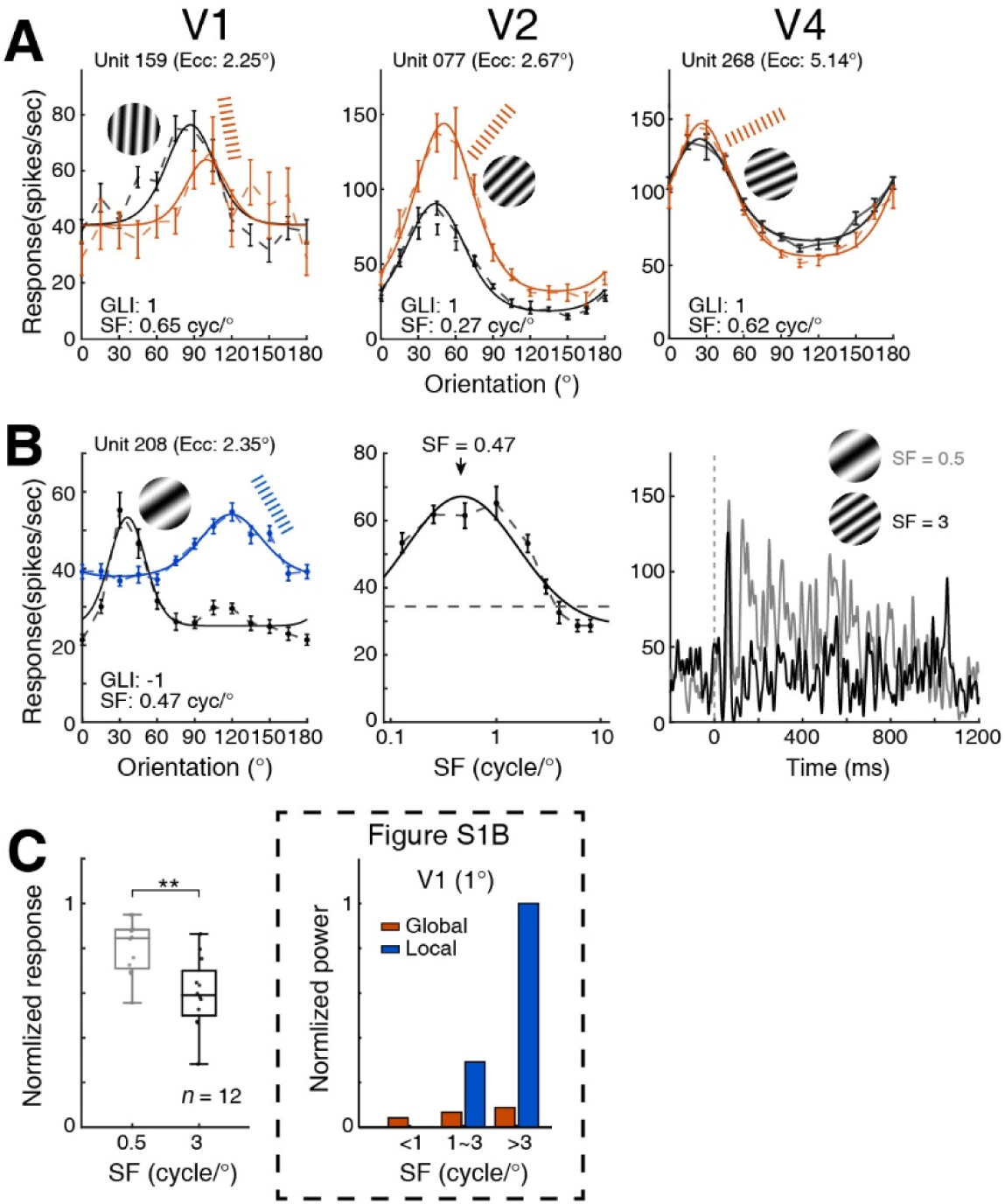
Neuronal responses in parafoveal regions across cortices from V1 to V4. **(A)** Orientation tuning curves of global-selective neurons with low-SF preferences in V1, V2, and V4. Stimulus icons indicate the preferred stimulus of each neuron. **(B)** A local-selective V1 neuron preferring low SF. Orientation tuning curves of the neuron are shown in the left panel. The middle panel shows its SF tuning curve with its average responses peaked at 0.47 cycles/°. The dashed line indicates baseline response. The right panel shows SDFs in response to grating stimuli with SFs of 0.5 and 3 cycles/°, respectively. Time 0 indicates stimulus onset. **(C)** Response comparison of local-selective V1 neurons when examined with SFs at 0.5 and 3 cycles/°, respectively. These neurons exhibited significant low-SF preferences to grating stimuli (\*\**p* < 0.01 with paired t-test). The dominance of the local-SF power in global–local stimuli accounts for the responses of these local-selective neurons (the inserted subfigure).

**Figure S6.**
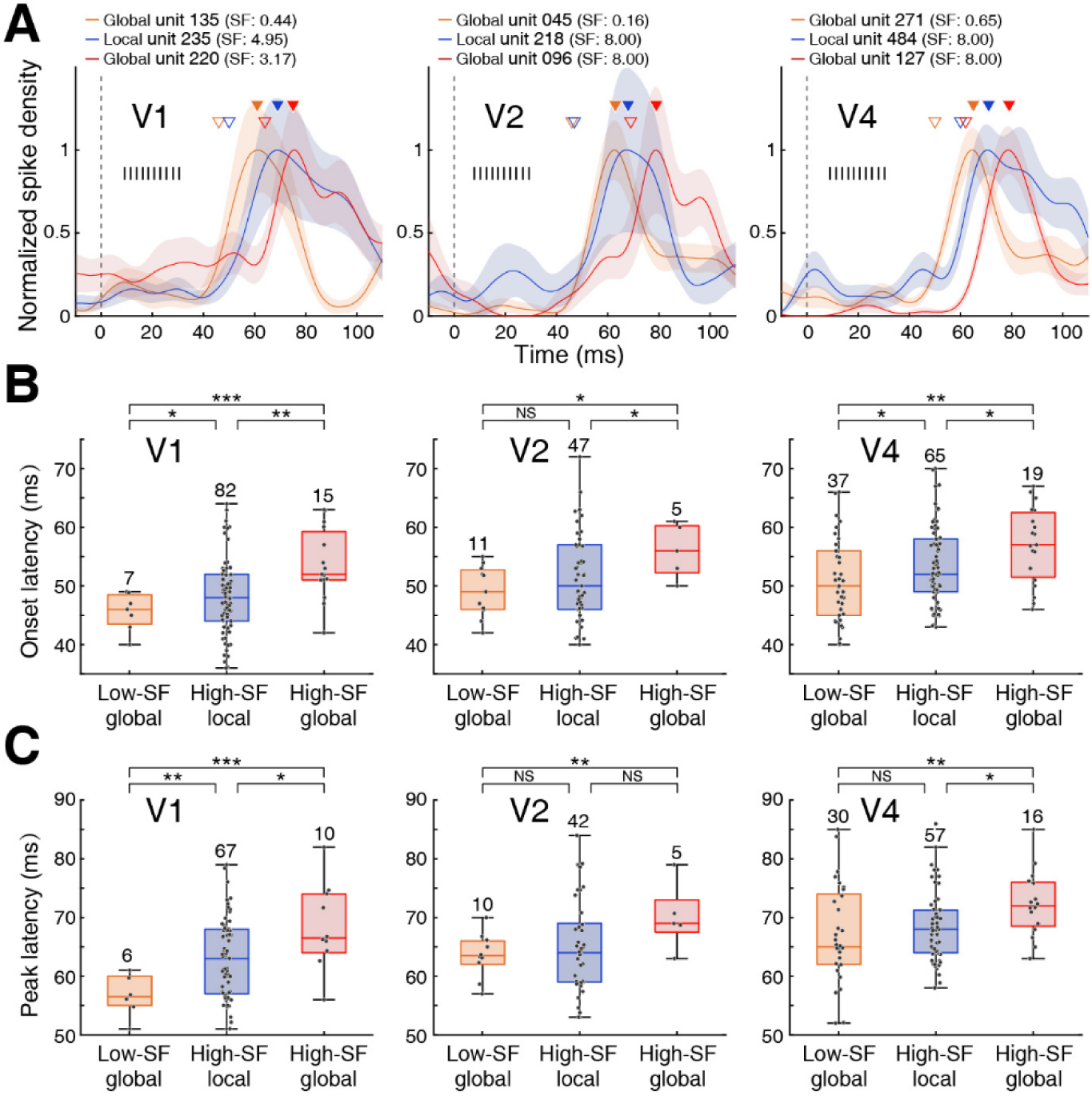
Response latencies to sinusoidal gratings and global–local stimuli from V1 to V4. **(A)** Temporal profiles of local and global responses of further individual neuron examples in V1, V2, and V4. Hollow and solid triangles indicate onset and peak latencies, respectively, for each neuron. **(B)** Comparison of onset latencies among the low-SF global, high-SF local and high-SF global groups within V1, V2, and V4 shown in main Figure 4F. The onset latencies (mean ± SEM) of low-SF global, high-SF local and high-SF global groups are 45.6 ± 1.2, 48.6 ± 0.8, and 53.7 ± 1.6 ms in V1, 49.1 ± 1.3, 51.4 ± 1.1, and 56.0 ± 2.1 ms in V2, 51.0 ± 1.2, 53.5 ± 0.8, and 57.1 ± 1.5 ms in V4, respectively. Whiskers indicate the minimum and maximum latencies; the central line denotes the median. \**p* < 0.05, \*\**p* < 0.01, \*\*\**p* < 0.001; NS, not significant. Numbers above the boxes indicate the number of neurons. **(C)** Comparison of peak latencies among the low-SF global, high-SF local and high-SF global groups within V1, V2, and V4 shown in main Figure 4F. The peak latencies (mean ± SEM) of low-SF global, high-SF local and high-SF global response groups are 56.7 ± 1.5, 63.1 ± 0.8, and 68.5 ± 2.3 ms in V1, 63.6 ± 1.2, 65.1 ± 1.2, and 70.2 ± 2.6 ms in V2, 67.3 ± 1.5,69.3 ± 1.1, and 72.4 ± 1.4 ms in V4, respectively. Regardless, the peak latency for high-SF global was the longest at each cortical area of V1, V2, and V4.

**Figure S7.**
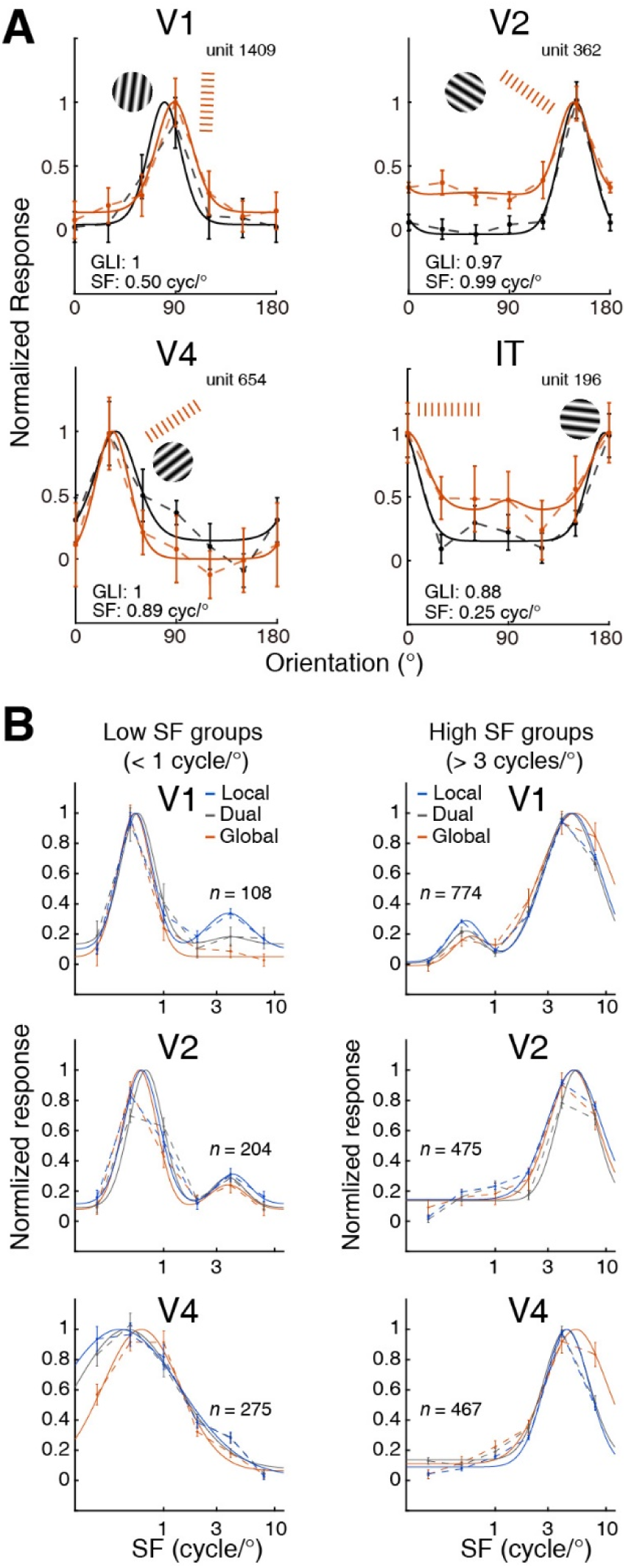
Neuronal responses to global–local stimuli in foveal regions across cortices. **(A)** Orientation tuning curves of global-selective neurons with low-SF preferences in V1, V2, V4, and IT. Stimulus icons indicate the preferred stimulus of each neuron. Neurons were recorded by two-photon imaging in foveal regions. **(B)** Averaged SF tuning curves of local-, dual-and global-selective neurons. Presented are SF tuning curves of neurons with preferred SFs below 1 cycle/° (the left column) and above 3 cycles/° (the right column) in V1, V2 and V4, respectively. Note that V1 and V2 neurons preferring low SFs also exhibited small peaks of high SF responses, whereas those preferred high SFs showing some low SF responses mainly in V1. For visualization, a dual-peaked log-Gaussian fitting was used. *n* indicates the number of neurons. Error bars represent SEM.

